# Ancestry-specific association mapping in admixed populations

**DOI:** 10.1101/014001

**Authors:** Line Skotte, Emil Jørsboe, Thorfinn Sand Korneliussen, Ida Moltke, Anders Albrechtsen

## Abstract

During the last decade genome–wide association studies have proven to be a powerful approach to identifying disease-causing variants. However, for admixed populations, most current methods for association testing are based on the assumption that the effect of a genetic variant is the same regardless of its ancestry. This is a reasonable assumption for a causal variant, but may not hold for the genetic variants that are tested in genome–wide association studies, which are usually not causal. The effects of non-causal genetic variants depend on how strongly their presence correlate with the presence of the causal variant, which may vary between ancestral populations because of different linkage disequilibrium patterns and allele frequencies.

Motivated by this, we here introduce a new statistical method for association testing in recently admixed populations, where the effect size is allowed to depend on the ancestry of a given allele. Our method does not rely on accurate inference of local ancestry, yet using simulations we show that in some scenarios it gives a dramatic increase in statistical power to detect associations. In addition, the method allows for testing for difference in effect size between ancestral populations, which can be used to help determine if a SNP is causal. We demonstrate the usefulness of the method on data from the Greenlandic population.

## Introduction

An individual’s risk of developing common complex diseases, such as type 2 diabetes, is influenced by specific genetic variants and identifying such variants using genome–wide association mapping studies (GWAS) has been a rapidly growing research field the last decade (Klein et al. 2005, Duerr et al. 2006, Burton et al. 2007, Unoki et al. 2008, Thorleifsson et al. 2009, Sparso et al. 2009, Holm et al. 2011).

So far, most GWAS have been performed in large homogeneous populations and this has led to important new findings (Morris et al. 2012, Voight B.F., et al. 2010, Scott et al. 2007). However, a few GWAS have now been performed in recently admixed populations (Moltke et al. 2014, The SIGMA Type 2 Diabetes Consortium 2013), and these studies have led to interesting new findings, suggesting that such studies can also be valuable to perform. This was recently shown very clearly in a study by Moltke et al. (2014) where a GWAS performed in the Greenlandic population, which has both Inuit and European ancestry, led to the identification of a variant that explains more than 10% of all cases of type 2 diabetes in Greenland. Notably, the variant had not been identified in earlier much larger studies of homogeneous populations like the European and East Asian populations, because it is very rare in these populations.

While being valuable, performing GWAS in recently admixed populations involves an important challenge: the admixture can bias the statistical test in the association mapping and lead to false discoveries. Statistical methods for association mapping that solve this challenge exist (Devlin & Roeder 1999, Price et al. 2006, Zhou & Stephens 2012), but these methods all share one essential limitation: they are based on the assumption that the tested genetic variant has the same effect regardless of which ancestral population it is inherited from. This assumption is reasonable for a disease-causing variant. However in GWAS, the disease-causing variant is often not tested directly. Instead a small fraction of common single nucleotide polymorphisms (SNPs) are genotyped and tested and the aim is to identify the subset of these SNPs, if any, that are indirectly associated with the disease, because they are located close to the causal SNP and therefore in linkage disequilibrium (LD) with it (see Figure 1). The effect size and strength of the association of a variant tested in a GWAS will therefore depend on the allele frequencies of the causal and tested variants and the strength of the LD between them. And importantly, since allele frequencies and LD patterns will often be different between different populations, this means that the effect size and the strength of association of a given variant may depend strongly on the ancestry of the chromosomal segment on which this variant is located. The extreme case shown in Figure 2A provides a simple illustrative example. Here the tested variant is present in both populations, but the causal variant is only present in ancestral population 1, where it is in complete LD with the tested variant. In this example, the tested variant is only associated with the phenotype when inherited from ancestral population 1 and thus the tested variant only has an effect in that population. This illustrates that the assumption of ancestry-independent effects, which most methods for association testing in admixed populations are based on, does not always hold in the context of GWAS in admixed populations. The example also illustrates another important point: the association with the tested variant in the example is weaker than the association with the causal variant, which in this case equals the association with the tested variant inherited from population 1. This means that a GWAS can potentially gain power by allowing for ancestry-specific effect sizes. Motivated by this, we here propose a statistical method for performing association mapping in admixed populations, named asaMap, that allows estimation and significance testing of ancestry-specific effect sizes. In individuals from admixed populations the local ancestry of an allele (corresponding to the red/blue color in Figure 2A) is not directly observable, but can sometimes be inferred (Patterson et al. 2004, Sankararaman et al. 2008, Price et al. 2009, Maples et al. 2013, Guan 2014). However, asaMap does not rely on accurately inferred local allelic ancestry because such inference can be prone to errors. Instead asaMap is based on a mixture model, where the mixture components are the phenotype distributions corresponding to given ancestries of the tested SNP and the mixture weights are the probabilities of these ancestries (for more details see Materials and Methods). This approach allows us to take the uncertainty of the ancestry of the individual alleles into account by allowing for all possible ancestries and weighting each possible ancestry according to its probability of being the true ancestry; the mixture weights. The mixture weights for a given SNP are in asaMap by default calculated from genome-wide admixture proportions, population specific allele frequencies and genotypes. However, asaMap also works with user provided mixture weights and thus allows users to use more complex models such as hidden Markov models (Patterson et al. 2004, Price et al. 2009, Guan 2014) for obtaining these probabilities. The mixture components in asaMap are based on a generalized linear model (GLM) framework. This has at least three advantages. First, it means we can correct for population structure by simply including principal components as covariates. Second, it makes asaMap very flexible, since it means that it – like a GLM – can be used to perform tests in a wide range of settings: asaMap can be applied to several different trait types (quantitative traits and case-control traits) as well as several different genetic effect types (additive and recessive effects). Third, it allows easy correction for any additional covariates such as sex or age.

**Figure 1:**
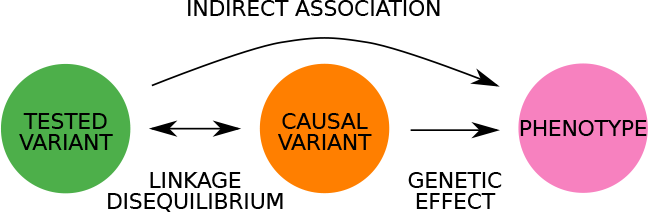
Indirect association between tested genetic variant and a phenotype.

**Figure 2:**
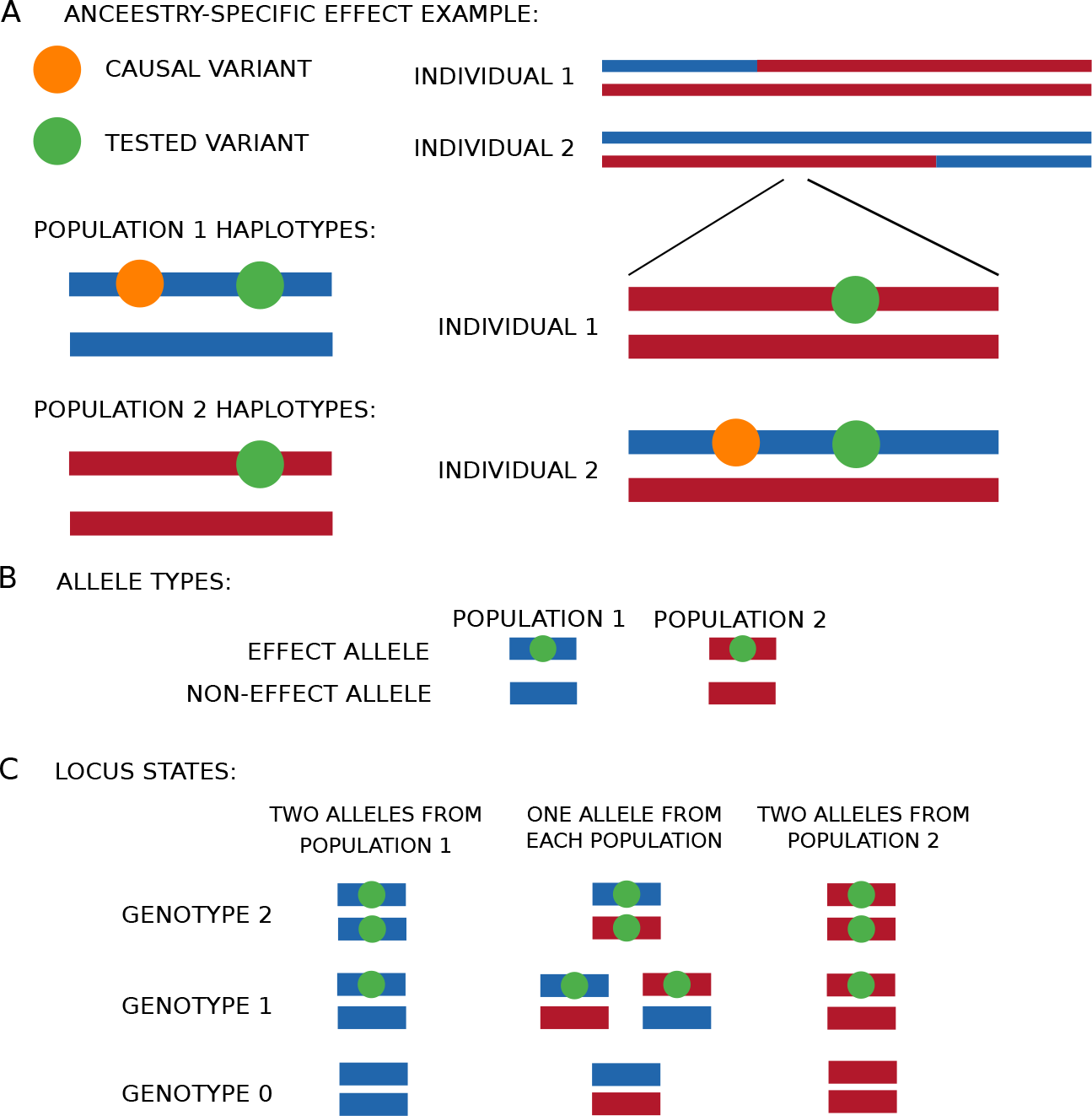
Ancestry-specific alleles. A: Extreme case of ancestry-specific effects in a population with ancestry from two populations: the tested variant is present in both ancestral populations, but the causal variant is only present in ancestral population 1, where it is in complete LD with the tested variant. The figure shows the homologous chromosomes of two admixed individuals with chromosomal segments colored according to which population the segments have been inherited from (population 1 is blue and population 2 is red). Both individuals carry one copy of the tested variant, however they have inherited them from different populations and only individual 2, who inherited the tested variant from population 1, carries the causal variant. B: The four ancestry-specific allele types that can occur at a diallelic autosomal locus in an individual from a population that consists of two ancestral admixing populations. C: The 10 distinguishable locus states (combinations of two ancestry-specific allele types) that can occur at a diallelic autosomal locus in an individual from a population that consists of two ancestral admixing populations.

We emphasize that asaMap is an association testing method and not a method for performing admixture mapping; a mapping approach where correlation between phenotype and inferred local ancestry is used to identify candidate regions (Patterson et al. 2004). asaMap is more similar to the methods of Pasaniuc et al. (2011), Yorgov et al. (2014) and Duan et al. (2017), where ancestry-specific effects are estimated and tested based on inferred knowledge about local ancestry. These methods focus on the potentially different effects of one specific allele (which we will refer to as “the assumed effect allele”) in the ancestral populations and they assume that the other allele (which we will refer to as “the assumed non-effect allele”) has the same mean phenotypic value in all ancestral populations after correcting for relevant covariates. Hence they ignore the possibility that it may be the assumed non-effect allele, and not the assumed effect allele, that mediates an ancestry-specific effect.

What makes asaMap differ from these methods is the fact that it does not require prior inferred knowledge of local ancestry, the fact that it enables correction for population structure, and that - although it also focuses on the potentially different effects of a specific assumed effect allele - it offers an additional test that allows the user to assess if it may be beneficial to use the other allele as the assumed effect allele instead.

In the following section, we describe the model behind asaMap in detail. Then using simulated data, we show that asaMap in some cases provides a substantial increase in power for association testing and that asaMap provides a framework that is even more flexible than the GLM, which is often used for association testing in GWAS. For example, we show that asaMap makes it possible to test whether a specific allele has different effects in different ancestral populations. Since it is reasonable to assume that a disease-causing allele has the same effect regardless of its ancestry, such a test can be used to reject that a SNP identified in a GWAS is causal assuming there is only one causal variant. Finally, using data from a GWAS in the admixed Greenlandic population (Moltke et al. 2014), we show that asaMap can provide increased power for SNPs that are in strong LD with the causal SNP and that asaMap can be used to discriminate between causal and non-causal variants, not only in simulated data, but also in real data.

## Materials and methods

### Model

Our asaMap model framework is based on a GLM. It therefore applies to both quantitative traits and case-control traits (i.e. dichotomous traits) and allows for both additive and recessive genetic effects. Here we describe the quantitative trait model for additive genetic effects, while detailed descriptions of the model for case-control data as well as recessive genetic effects can be found in the Supporting Information.

As argued in the introduction, the strength of an association between genotype and phenotype is likely to depend on which of the ancestral populations a genetic allele has been inherited from. Thus, instead of estimating a single genetic effect of a given allele, we here allow for population-specific genetic effect sizes, each of which we denote *β*_*k*_ for an allele inherited from ancestral population *k*. Below we describe the model that allows us to do this. The description is limited to two ancestral populations, but could be extended to more than two populations.

#### Mixture model

We assume that we are analyzing data from *N* individuals from an admixed population that is a mixture of two ancestral populations. An individual from such a population will at any given diallelic autosomal locus have inherited each of its two chromosome copies from one of the two ancestral populations, and each of these will either carry the effect allele or not. Thus there are four possible ancestry-specific allele types, as illustrated in Figure 2B. As a consequence, each individual will have one of 16 different ancestry-specific allele type combinations at any given diallelic autosomal locus. We will refer to these ancestry-specific allele type combinations as *locus states*, *s*. The 10 distinguishable locus states are shown in Figure 2C. Since the true locus state is not observable from genotype data, and since it is challenging to infer we chose a model that allows us to take the uncertainty of this state into account. This model is based on the observation that when all that can be observed are the genotypes, i.e. the total number of copies of a specified allele present at the tested locus in each individual, the likelihood function for the measured phenotypes, *Y* = (*y*_1_, *y*_2_, …, *y*_*N*_), takes the form

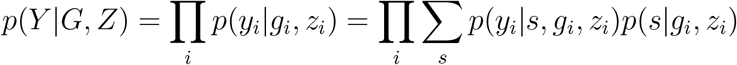

where *G* = (*g*_1_, *g*_2_, …, *g*_*N*_) is a vector of all observed genotypes at the tested locus, *Z* is a matrix of appropriately chosen covariates, *z*_*i*_ is the vector of covariate values for individual *i*, the product runs over all individuals *i* = 1 … *N* and the sum runs over all possible locus states *s*. Assuming that the trait is conditionally independent of the observed genotypes *G* given the latent variable *s* and the covariates *Z*, this likelihood also takes the form

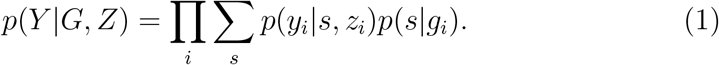

This means that we can model the probability of the measured phenotypes *Y* as a mixture of phenotype distributions, where each mixture component is the phenotype distribution given a specific locus state *s*, *p*(*y*|*s*, *z*), and the corresponding mixture weight is the probability *p*(*s*|*g*) of that locus state given the observed genotype. Notably, this modeling approach makes it very easy to take the uncertainty of the unobserved ancestry into account, since this uncertainty is explicitly included in the model in the form of the mixture weights. Furthermore, the above likelihood function is a function of our parameters of interest, namely the population-specific effects *β*_*k*_, via the mixture components *p*(*y*|*s*, *z*). Hence we can use the model both for estimating the ancestry-specific effects *β*_*k*_ and for performing association testing. Specifically, in asaMap the *β*_*k*_s are estimated using maximum likelihood estimation and testing for association is performed using likelihood ratio tests. Below is a detailed description of how we model the mixture components and the mixture weights in the likelihood function, followed by a detailed description of the parameter estimation and testing procedures.

#### Mixture components

For a given quantitative trait, *Y*, the mixture component, i.e. the phenotype distribution *p*(*y*|*s*, *z*), is based on a linear regression model. We assume that given the locus state, *s*, the phenotype *y*_*i*_ for a single individual *i* follows a normal distribution with mean given by the linear predictor

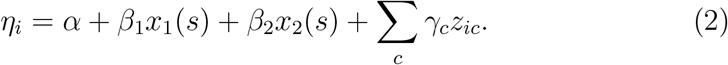

Here *α* parameterizes the intercept (baseline) and each additional covariate *c*, such as principal components for correcting for potential confounding by population structure (Price et al. 2006), enters the model in the *z*_*ic*_ with effects γ_*c*_. Finally, assuming an additive model, *x*_1_(*s*) and *x*_2_(*s*) are the counts of the assumed effect allele from population 1 and 2, respectively, for a locus in locus state *s*. If one instead assumes a recessive model, the definition of the terms *x*_1_(*s*) and *x*_2_(*s*) is different and is described in more details in the Supporting Information.

#### Mixture weights

The simplest approach to calculating the mixture weights, i.e. the probabilities of the different possible locus states for an individual given its genotype *g*, is to use the genome-wide admixture proportions *q* = (*q*_1_, *q*_2_) and the population specific allele frequencies *f*, both of which can be inferred using standard software tools such as ADMIXTURE (Alexander et al. 2009). Note that for convenience we have excluded the subscript *i* here. In the case of two admixing populations the genome-wide admixture proportions are *q* = (*q*_1_, *q*_2_), where *q*_1_ and *q*_2_ is the fraction of its genome that has been inherited from population 1 and 2 respectively, and the population specific allele frequencies are *f* = (*f*_1_, *f*_2_), where *f*_1_ and *f*_2_ is the frequency of assumed effect allele in population 1 and 2, respectively.

We can use *q* and *f* to calculate the probability of the locus state *s* given a genotype *g* in three steps after introducing the notation *s* = (*a*, *t*), where *a* = (*a*_1_, *a*_2_) is the ordered allelic ancestry and *t* = (*t*_1_, *t*_2_) is the ordered allelic genotype (with *t*_1_ + *t*_2_ = *g*). In the first step we consider the conditional distribution of the ordered allelic genotype *t* = (*t*_1_, *t*_2_) given genotype *g* = *t*_1_ + *t*_2_, which takes the form:

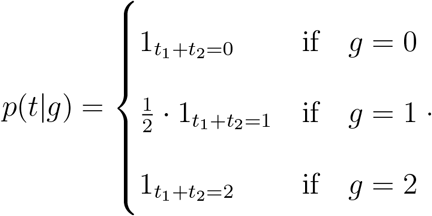

In the the second step we consider the probability of the ordered allelic ancestry *a* = (*a*_1_, *a*_2_) given the ordered allelic genotype *t* = (*t*_1_, *t*_2_). Here we use the genome-wide admixture proportions *q* to give the probability 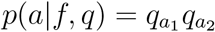 of ancestry *a* assuming independent ancestry of alleles and we use the corresponding population specific allele frequencies *f* to calculate the probability of ordered allelic genotype given the ordered allelic ancestry 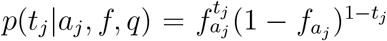. With this the desired probability of the ordered allelic ancestry *a* given the ordered allelic genotype *t* can be calculated by

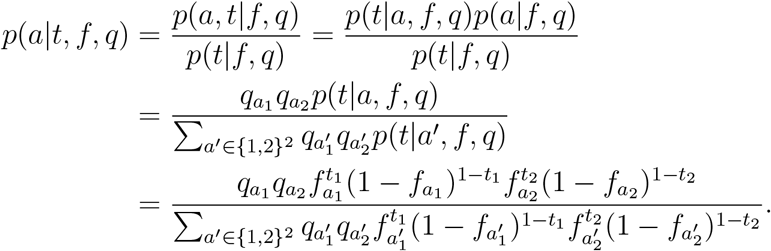

The third step combines the results of the first two steps to calculate the conditional distribution of locus states given the observed genotype using *q* and *f*, splitting the joint conditional probability into the two terms we have just derived:

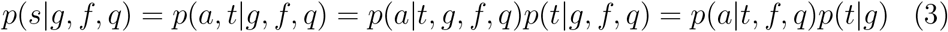

In the last derivation step we exploit the fact that *a* depends on *g* via *t*, and that *t* only depends on *g*. This three-step procedure is the default procedure used in asaMap. However, the user can also supply the distribution across locus states to asaMap and can therefore choose to for instance calculate it based on the output from local ancestry inference software (such as (Patterson et al. 2004, Price et al. 2009, Guan 2014)) instead.

### Parameter estimation and hypothesis testing

#### Parameter estimation

The parameters of the model described above (and of the sub-models relevant for testing purposes described below) are estimated using maximum likelihood based on the likelihood function given in equation 1 (using the details provided in equation 2 and equation 3). Optimization of this likelihood function must be done numerically and we have developed an expectation-maximization (EM) algorithm for this (see Supporting Information for details) that provides faster convergence than standard all-purpose numerical optimization algorithms such as BFGS.

#### Hypothesis testing

Standard GLM based methods that are commonly used for association testing typically assume an additive effect and make use of statistical tests comparing two models: a model where a given allele has a genetic effect versus a nested model where it has no effect. In asaMap we allow the effect sizes to be specific to ancestral populations and therefore several more nested models can be compared. Specifically, for an additive genetic effect, then assuming two admixing populations five models M1-M5 are available for comparison. The full model M1 allows separate genetic effects for each of the two ancestral population: *β*_1_ and *β*_2_ (see equation 2). The sub-model M2 assumes no effect in population 1 (i.e. *β*_1_ = 0). The submodel M3 assumes no effect in population 2 (i.e. *β*_2_ = 0). The sub-model M4 assumes that the effect sizes are the same in both ancestral populations (i.e. *β*_1_ = *β*_2_), and finally the null sub-model M5 assumes that there is no effect in any of the populations (i.e. *β*_1_ = *β*_2_ = 0). An overview of these additive models is given in Table 1 and a detailed description can be found in the Supporting Information. For recessive genetic effects the standard GLM based methods for association mapping tests a model where carrying two copies of a given allele has an effect on the individual’s phenotype versus a model where carrying two copies of the allele has no effect. Here asaMap allows the effect size to be specific to the ancestry combination of the two allele copies. To do this seven (sub-)models R1-R7 described in the Supporting Information and Table S1 are implemented.

**Table 1:** Description of the five different possible additive models when there are two ancestral populations. *β*_1_ is the effect size of alleles from population 1 and *β*_2_ is the effect size of alleles from population 2. Comparing different pairs of nested models leads to the tests described in Table 2. For a description of recessive models, see Table S1.

| Model | Hypothesis | The model assumes |
| --- | --- | --- |
| M1 | $(\beta_1, \beta_2) \in R^2$ | Population specific effects |
| M2 | $\beta_1 = 0, \beta_2 \in R$ | No effect in population 1 |
| M3 | $\beta_1 \in R, \beta_2 = 0$ | No effect in population 2 |
| M4 | $\beta_1 = \beta_2 \in R$ | Same effect in both populations |
| M5 | $\beta_1 = \beta_2 = 0$ | No effect in any population |

In asaMap hypothesis tests regarding the ancestry-specific effect sizes is carried out using likelihood ratio tests comparing nested models among the models just described. The implemented models under the additive effect assumption allow us to test if there is an effect in any population (M1 vs. M5), an effect in population 1 (M3 vs. M5), an effect in population 2 (M2 vs. M5), and a difference in the effect specific to the two ancestral populations (M1 vs. M4). Furthermore, they allow us to test if there is an effect assuming that it is the same in the two populations (M4 vs. M5). Note that this latter test is equivalent of the standard test for association performed using a GLM and it has been implemented in asaMap to enable comparison of the other tests to the commonly used standard GLM based testing approach. In addition, the tests M1 vs. M2 and M1 vs. M3 are also implemented and may be used for performing model checks of the tests based on models M2 and M3. An overview of the implemented tests for additive genetic effects is given in Table 2. The corresponding tests comparing nested recessive models are described in the Supporting Information and Table S2.

**Table 2:** Description of the possible tests comparing different nested models (described in table 1) assuming an additive genetic model.

| Models | Tests if there is |
| --- | --- |
| M1 vs. M5 | an effect in any population |
| M1 vs. M2 | an effect in population 1 |
| M1 vs. M3 | an effect in population 2 |
| M1 vs. M4 | a different effect in the two populations |
| M2 vs. M5 | an effect in population 2 assuming no effect in population 1 |
| M3 vs. M5 | an effect in population 1 assuming no effect in population 2 |
| M4 vs. M5 | an effect assuming it is the same in both populations |

#### Implementation

The estimation and testing procedures described above have been implemented in the software asaMap available at https://github.com/e-jorsboe/asaMap (C++) and https://github.com/lineskotte/asamap (R-package), making the method applicable to large scale genome wide association studies.

### Testing if the assumed non-effect allele has a population specific effect

Like similar methods by Pasaniuc et al. (2011), Yorgov et al. (2014) and Duan et al. (2017), asaMap focuses only on one specific allele, the assumed effect allele, when allowing for ancestry specific effects, as illustrated in Figure S1A. Thus it ignores the possibility that the other allele, the assumed non-effect allele, could mediate an ancestry-specific effect (Figure S1B). To test if this is the case, and thus that association testing focused on this other allele instead may be preferred, we introduce an additional model M0 (Figure S1C), which is an extension of model M1.

In M0 the phenotype *y*_*i*_ for a single individual *i* follows a normal distribution with mean given by the linear predictor:

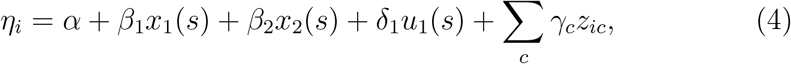

Hence M0 is M1 extended with an additional predictor *u*_1_, which is the count of the assumed non-effect allele from population 1, and the additional parameter *δ*_1_, which is effect size of the assumed non-effect allele in population 1 (see Figure S1C). The other symbols are the same as in equation 2. We use model M0 by comparing it to model M1 (Table 1) and testing if we can reject the null hypothesis of *δ*_1_=0. If this null hypothesis can be rejected, this provides evidence that the assumed non-effect allele has a population specific effect and that this allele may therefore be beneficial to use as the assumed effect allele instead.

The above test is for additive effects. We propose a similar test for recessive effects, in which an additional model, R0, is compared to model R1. For details about R0 and this test, see Supporting Information.

### Comparison between asaMap and the LAAA test

As previously mentioned Duan et al. (2017) has proposed a test similar in idea to asaMap. It is called “LAAA” and it is equivalent to comparing models M0 and M5 in asaMap, except that LAAA does not account for uncertainty in the ancestry estimates. This means that the inclusion of M0 in asaMap allows us to compare the power between asaMap and LAAA by simply comparing different tests in asaMap. Specifically, we performed such a power comparison by comparing the power of the M1 vs. M5 test and M0 vs. M5 test in asaMap.

### Correcting for population structure

To correct for population structure in the real data (described below), we include as covariates the first 10 principal components calculated from a genotype-based covariance matrix (Price et al. 2006). We are aware that a more powerful approach would be a mixed effects model approach similar to (Kang et al. 2008) or (Zhou & Stephens 2012), but we have not succeeded in implementing this in a computationally tractable way in the context of the models proposed here due to the sum across locus states. Since it has been shown that including the first 10 principal components does not always fully correct for population structure, we highly recommend to make QQ-plots to assess the outcome of asaMap - like in all other genetic association studies. If an inflation of the test statistics is observed one potential solution is to use genomic control (Devlin & Roeder 1999).

### Simulated data

To be able to assess asaMap, we carried out analyses of simulated samples with genetic ancestry from two admixing populations. We simulated data from a total of nine scenarios. In each of these scenarios we simulated data from a SNP locus, which is assumed not to be causal, but to be in LD with a causal variant. For all nine scenarios we simulated data from a total of 2500 individuals with admixture proportions from population 1, *q*_1_, in the set {0, 0.25, 0.5, 0.75, 1} (500 individuals for each value). What varies between the scenarios is the frequency of the tested variant in the two populations, *f* = (*f*_1_, *f*_2_), the effect sizes in the two populations (*β*_1_ and *β*_2_), the type of trait (quantitative or case-control) and the underlying genetic effect model (additive or recessive). For a description of the nine scenarios see Table 3.

For all scenarios, we followed the same simulation procedure: for each individual we sampled the ordered allelic ancestry *a* = (*a*_1_, *a*_2_) based on the individual admixture proportions *q*_*i*1_. Then based on this ancestry *a* and the allele frequencies *f* in the ancestral populations, we sampled the ordered allele types *t* = (*t*_1_, *t*_2_) for each individual. Knowing *a* and *t*, the genotype and locus state for each individual is known and based on the latter the phenotype of each individual is simulated using the relevant phenotype distribution (quantitative or case-control), the relevant genetic model (additive or recessive) and scenario specific effect sizes (*β*_1_ and *β*_2_). For quantitative traits we generated the phenotype value using a normal distribution with variance 1 and for case-control studies we generated the disease status using a binomial distribution.

These simulations gave us access to association data where the true ancestry-specific effects are known and therefore allowed us to assess the consistency and unbiasedness of the estimators in asaMap. The simulations also allowed us to assess the power of the tests for association implemented in asaMap. Furthermore, because we explicitly simulate the ancestry-specific allele type combinations (locus states), the simulations allowed us to compare the power of the tests in asaMap to the hypothetical power of a test where the true locus states, which in reality are unobservable, are known.

To be able to assess the impact of the choice of admixture proportions, we also performed the same simulations with admixture proportions estimated from a real dataset. Here we used the admixture proportions from Inuit Health in Transition (IHIT) cohort of admixed Greenlandic individuals estimated in Moltke et al. (2014). We note that the two ancestral populations are not equally represented in these individuals, and we therefore repeated the simulations with the admixture proportions for population 1 and 2 interchanged.

**Table 3:** Simulated scenarios. All scenarios have 2500 individuals and individual admixture proportions from population 1 in {0, 0.25, 0.5, 0.75, 1}. The allele frequencies *f* = (*f*_1_, *f*_2_) in the two ancestral populations are shown in the “Frequencies” column. For each scenario we vary the effect size, however, they are restricted in some scenarios as shown in the “Effects” column. E.g. in scenario A1 there is only an effect in population 1, so here the effect size in population 2 is restricted to 0. When there is an effect in both populations, we assume that they are the same. The type of trait simulated is shown in the “Trait” column and the simulated effect model is shown in the “Model” column.

| Scenario | Frequencies, $f=(f_1, f_2)$ | Effects | Trait | Model |
| --- | --- | --- | --- | --- |
| A1 | (0.1, 0.3) | Population 1 | Quantitative | Additive |
| A2 | (0.1, 0.3) | Both | Quantitative | Additive |
| A3 | (0.1, 0.3) | Population 2 | Quantitative | Additive |
| B1 | (0.2, 0.2) | Population 1 | Quantitative | Additive |
| B2 | (0.2, 0.2) | Both | Quantitative | Additive |
| B3 | (0.2, 0.2) | Population 1 | Quantitative | Recessive |
| C1 | (0.2, 0.2) | Population 1 | Case-Control | Additive |
| C2 | (0.2, 0.2) | Both | Case-Control | Additive |
| C3 | (0.4, 0.4) | Population 1 | Case-Control | Recessive |

### Data for *TBC1D4* in a Greenlandic cohort

To illustrate that allowing for ancestry-specific effect sizes can be informative and appropriate for real data we applied asaMap to genotype data in combination with measurements of 2-hour (2-h) plasma glucose levels of 2575 individuals in the IHIT cohort from the admixed Greenlandic population (Moltke et al. 2014, Jorgensen et al. 2013). More specifically, we applied asaMap to five genotyped SNPs in the *TBC1D4* gene (rs61736969, rs7330796, rs1062087, rs2297206 and rs77685055). In Moltke et al. (2014) all five SNPs were found to be strongly associated with an increase in 2-h plasma glucose levels. rs7330796 was the lead SNP in the discovery part of the study, which was based on SNP chip data, and rs61736969 is the causal SNP that was in a later part of the study identified from sequencing data. The three remaining SNPs were also identified from sequencing data in the search for the causal variant.

The study received ethics approval from the Commission for Scientific Research in Greenland (project 2011-13, ref. no. 2011-056978; and project 2013-13, ref.no. 2013-090702) and was conducted in accordance with the ethical standards of the Declaration of Helsinki, second revision. Participants gave written consent after being informed about the study orally and in writing.

## Results

To investigate the cost and benefits of asaMap compared to a standard GLM based test we first applied both methods to simulated data to compare their statistical power and to assess important statistical properties of asaMap, like bias and false positive rates. Using the same simulated data we also compared asaMap to another similar test, the LAAA test recently proposed by Duan et al. (2017). Finally, we applied both the standard GLM based methods to real data to compare the range of their potential usage. In all cases we investigated populations that are mixtures between two populations.

Note that because the effect sizes are allowed to be specific to ancestral populations numerous models can be compared in asaMap (for an overview see Tables 1 and 2 for additive genetic effect based models and Tables S1 and S2 for recessive genetic effect based models). Note also that the test in asaMap that compares models M4 and M5 (R6 vs. R7) is equivalent of the standard GLM based test for association. In the following we will therefore perform the comparison of the standard GLM based test and asaMap by comparing the M4 vs. M5 (R6 vs. R7) test with a range of other tests available in asaMap.

### Simulation-based results

To assess the power and other statistical properties of asaMap we first simulated data for individuals with genetic ancestry from two admixing populations according to nine scenarios (scenarios A1-C3); six with quantitative traits and three with case-control traits (Table 3, see Materials and Methods for details).

#### Power assessment for quantitative traits

First we simulated a scenario (scenario A1) with a causal variant that is only present in one of the ancestral populations, but with the tested variant present in both ancestral populations, causing the tested variant to have population-specific effects. More specifically, the tested variant was simulated to have an additive effect, with an effect size in population 1, *β*_1_, that varied in the range [0, 1.5], and with no effect in population 2 since the causal variant is not present in this population. Furthermore, the tested variant was simulated to have a frequency 10% in ancestral population 1 and 30% in ancestral population 2. When applying asaMap to data from this scenario, the standard GLM based (linear regression) test (M4 vs. M5), where the effect size is assumed to be the same in both populations, required much larger effect sizes for full statistical power than the full test where the effect size is not assumed to be the same (M1 vs. M5), see Figure 3:A1. Also, the test of whether there is an effect in population 1 (M1 vs. M2 and M3 vs. M5) was slightly more powerful than the full test (M1 vs. M5), which is expected since it has only 1 degree of freedom and the full test has 2 degrees of freedom. Finally, asaMap could test whether there is a difference in the effect sizes between ancestral populations (M1 vs. M4) with good statistical power, even for variants with effect sizes lower that those detectable using the standard GLM based test.

Second, we simulated a scenario (A2), where the tested variant has the same effect in both populations, which corresponds to a situation in which the causal allele is tested. In this scenario, the test of M1 vs. M5 was less powerful than the standard GLM based test (M4 vs. M5), which was expected because of the extra degree of freedom (scenario A2, Figure 3). However, interestingly the difference in power is very small. On the other hand, the test, M3 vs. M5, of whether there is an effect in population 1, where the tested variant has the lowest frequency is markedly less powerful. Notably, in this scenario we do not reject that the effect is the same for the two populations (M1 vs M4).

Third, we simulated a scenario (A3) in which the tested variant only has an effect in population 2, where the variant allele occurs with the highest frequency. In this scenario, the statistical power of the test, M1 vs. M4, of whether there is a difference in effect sizes, was the same as for scenario A1, but, unlike in scenario A1 the standard GLM based test was almost as powerful as the remaining tests (scenario A3, Figure 3).

Next, we simulated a scenario (B1) where the tested variant has the same frequency of 20% in both ancestral populations. As was the case for scenario A1, the test with the best statistical power was the test of whether there is an effect in population 1 (M3 vs. M5) (scenario B1, Figure 3). Notably, this test was only slightly more powerful than the full test for effects (M1 vs. M5), but both these test provided remarkable improvements in power compared to the standard GLM based test of effect assuming same effect in both populations (M4 vs. M5).

**Figure 3:**
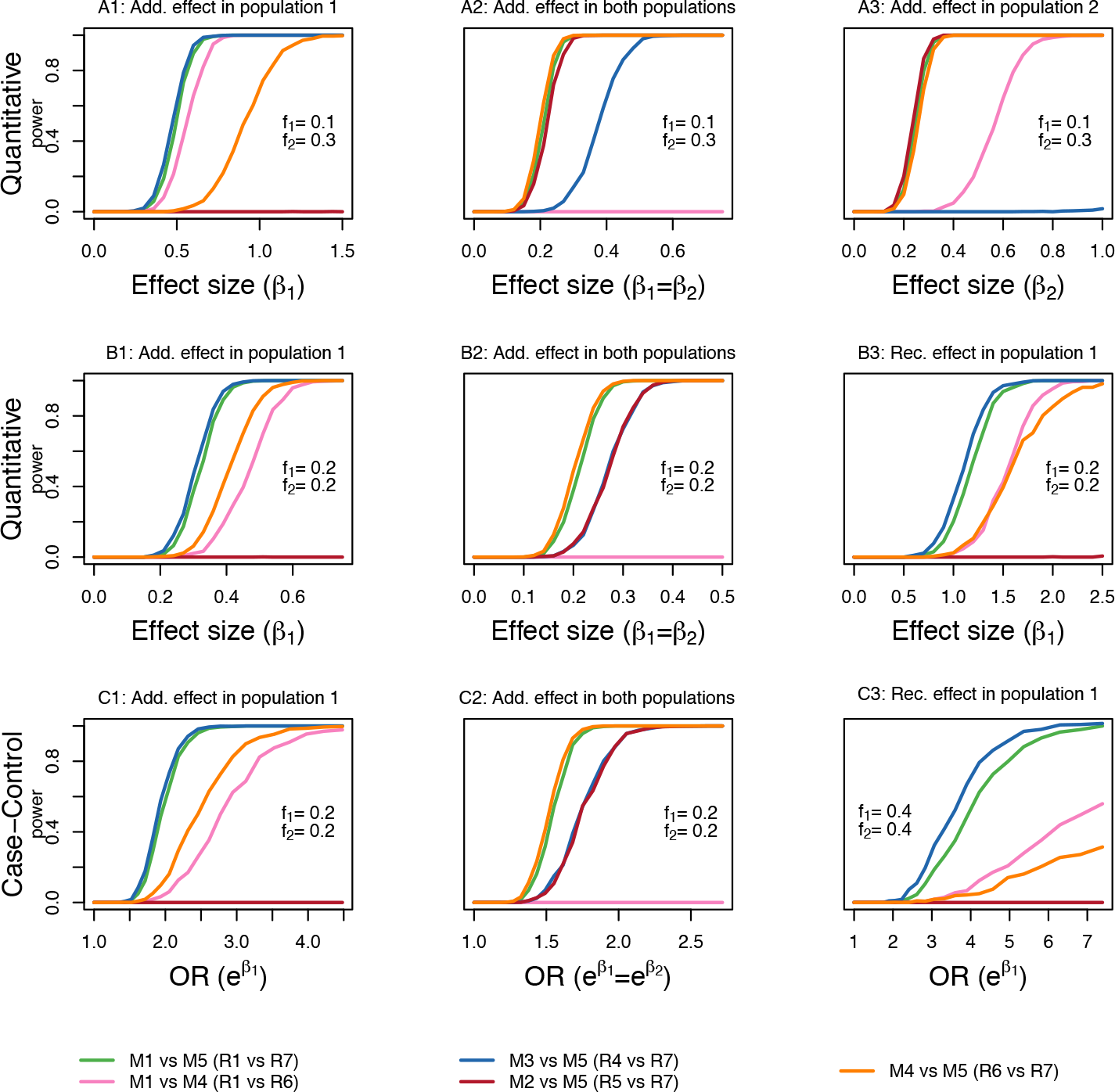
Power assessment results obtained by applying asaMap to data simulated from nine different. The curves show the fraction of simulated p-values that are smaller than 10^−8^, based on 1000 simulations for each effect size for each scenario. The simulated population specific effects *β*_1_*, β*_2_ are additive, except for scenario B3 and C3. *f*_1_ and *f*_2_ denote the population allele frequencies.

We then simulated scenario B2, which was identical to scenario B1 with the exception that the tested variant was simulated to have the same effect in both ancestral populations. Again, as was the case for scenario A2, the expected loss of power of M1 vs. M5 due to the more complicated modeling compared to M4 vs. M5 was very small (scenario B2, Figure 3), while the single population tests (M1 vs. M2 and M1 vs. M3) were less powerful.

Finally, to compare the power of the different tests for a variant with a recessive effect, we simulated a scenario (scenario B3), where the frequency of the tested variant was 20% in both ancestral populations and it had a recessive effect in population 1 and no effect in population 2. The results for this scenario were similar to the results for scenario A1 and B1: the test of an effect in population 1 (R4 vs. R7) was slightly more powerful than the full test (R1 vs. R7) and both represent remarkable improvements compared to the standard GLM based test (R6 vs. R7) (Figure 3).

#### Power assessment for case-control study data

For case-control study data we simulated from a population of mixed ancestry where the tested variant has a frequency of 20% in both populations. The effect of the allele was simulated as log-additive in the logistic model and either only present in ancestral population 1 (scenario C1, Figure 3) or present in both ancestral populations (scenario C2, Figure 3). The results are similar to the results for the quantitative trait versions of these scenarios (scenarios B1 and B2). In scenario C1 the asaMap test for an effect in ancestral population 1 (M3 vs M5) is the most powerful test, slightly more powerful than the asaMap test for an effect in any population (M1 vs. M5), and both these tests outperform the standard GLM based (logistic regression) test for association. In scenario C2, where the effect is present in both ancestral populations, the standard GLM based test is as expected the most powerful, but only slightly better than the asaMap test for an effect in any population (M1 vs. M5).

We also simulated a similar case-control scenario (scenario C3), where the effect of the tested variant is recessive and present only in ancestral population 1 (scenario C3, Figure 3). Note that to reach any statistical power for the simulated odds ratios (ORs), we here allowed the frequency of the tested allele to be 40% in both ancestral populations. In this scenario, the standard GLM based test of whether there is an effect assuming it is the same in both populations (R6 vs. R7), does not reach satisfactory statistical power for any realistic ORs. The tests that allow for population-specific effects (R1 vs. R7 and R4 vs. R7) on the other hand perform much better, although they also require quite high ORs to reach full statistical power.

#### Power assessment with asymmetrical admixture proportions

We based the above simulations on a set of artificially chosen admixture proportions with a symmetrical contribution from the two ancestral populations. To investigate the impact of different admixture proportion distributions on the statistical power of asaMap, we performed additional power simulations using the same 9 scenarios, but different of admixture proportions. In particular, we used admixture proportions inferred from a real population cohort with two ancestral populations of which one has contributed markedly more genetically than the other to the cohort (ca. 25% vs. 75%, see Moltke et al. (2014)). We performed the additional simulations both with population 1 being the population that has contributed the most (Figure S6) and with population 1 being the population that has contributed the least (Figure S7). The results show that when the ancestry specific effect lies in the population that has contributed the most, the standard GLM based test (M4 vs. M5) has power similar to M1 vs. M5 and M2 vs. M5 (see scenario A1, B1, B3, C1 and C3 in Figure S6). In contrast, when the ancestry specific effect lies in the smaller population, the standard GLM based test (M4 vs. M5) has less power than M1 vs. M5 and M2 vs. M5 (see scenario A1, B1, B3, C1 and C3 in Figure S7). Hence the admixture proportions have a marked impact on how much power there is to gain from using asaMap compared to the standard GLM based test. And notably the gain is largest when the ancestry specific effect is in the least contributing ancestral population.

#### Bias, consistency and false positive rates

Besides using the simulated data for power comparisons, we also used it for assessing asaMap’s estimators for population specific effects. We did this for all nine simulated scenarios (A1-3, B1-3 and C1-3). This showed that asaMap’s estimators are unbiased for these scenarios (Figure S2). For all simulation setups, we also simulated data under the null, i.e. without any effect in any of the populations, and applied all tests available in asaMap to the data to assess if asaMap has a controlled false positive rate. This was done to ensure that the uncertainty in ancestry does not lead to inflated test statistics. More specifically, we did this for the shared null model of scenarios A1-A3, the shared null of scenarios B1-B2, the null of scenario B3, the shared null of scenario C1-C2 and the null of scenario C3, which means that we performed the assessment both in the context of quantitative traits and of case-control data and both for variants with additive effects and variants with recessive effects. The corresponding QQ-plots of the p-values achieved show that the false positive rate is indeed controlled for in all the tests (Figure S3).

**Figure 4:**
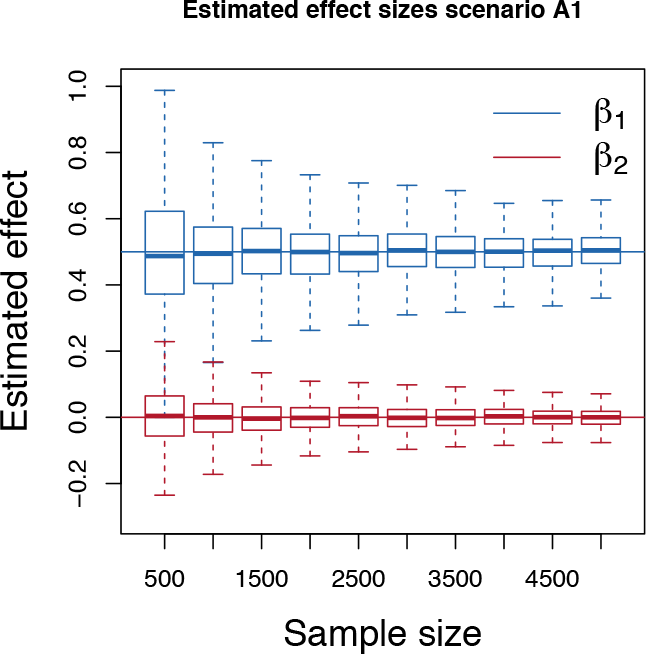
Consistency results obtained by applying asaMap to data from scenario A1 simulated with a fixed effect size of 0.5 in population 1 and 0 in population 2 and with sample sizes ranging from 500 to 5000. Each box corresponds to 1000 simulations.

Finally, to assess consistency of the estimators, we next re-simulated scenario A1 with increasing sample sizes and a fixed population specific effect size of 0.5 in population 1 and 0 in population 2. This showed that asaMap’s estimators are consistent and that the decrease in variance with increasing sample size is consistent with the expected 1/*n* relation (Figure 4).

#### Assessment of potential power gain from knowing the locus states

In the process of simulating ancestry-specific association data for the above power assessment, we explicitly simulated the ancestry-specific allele type combinations (locus states, *s*), which are not directly observable in real data. This allowed us to compare the tests implemented in asaMap with hypothetical tests of equivalent models based on known locus states. As expected, the tests based on correctly known locus states is more powerful and the variance of the estimators is a bit smaller (Figure 5, results only shown for scenario A1), which shows that if the ancestry of each allele copy was known without error there is potential for an even larger increase in statistical power.

#### Assessment of the test for whether the assumed non-effect allele has an ancestry-specific effect

In the above assessments, we used the simulated effect allele as the assumed effect allele in all cases. To assess how well our proposed tests for whether the assumed non-effect allele has an ancestry-specific effect and thus should be used as the assumed effect allele, we again simulated data from the nine scenarios from Table 3. However, this time we performed the asaMap analyses of the data with the simulated effect allele as the assumed non-effect allele in statistical models M1 (R1).

**Figure 5:**
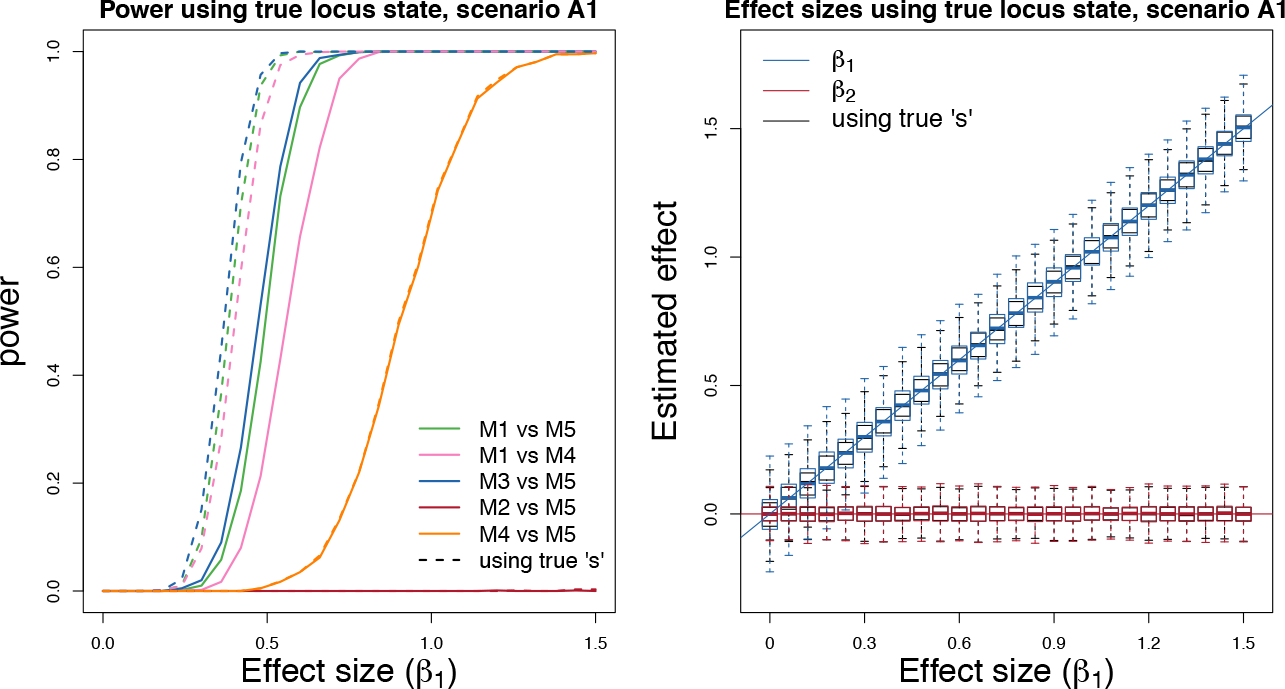
Comparison of power and bias results obtained by applying asaMap with or without known locus state (*s*) to simulated data from scenario A1. Left: Results of power comparison. The power obtained with the true (simulated) locus states are shown with dashed curves and the power obtained without are shown with solid curves. Right: Results of bias comparison. Box plots of estimates obtained with the true (simulated) locus states are shown in black, whereas estimates obtained without are shown in colors.

We compared M0 vs. M1 (R0 vs. R1) to see if these tests correctly reject that *δ*_1_ = 0 (equation 4) and that *δ*_1_ = *δ*_2_ = 0 (equation 7) when the effect allele is misspecified. The tests have high power to correctly reject the choice of effect allele (Figure S4) in scenarios A1, A3, B1, B3, C1 and C3. The simulated effect is not population-specific in scenarios A2, B2 and C2 and the test is correct not to reject the choice of effect allele here. To check that the false positive rate of the test for a misspecified effect allele is controlled, we also applied the M0 vs. M1 (R0 vs. R1) test with the simulated true effect allele as the assumed effect allele in the statistical model M1 (R1). Here, by simulation *δ*_1_ = 0 in the statistical model M0 and the corresponding hypothesis should only be rejected at rate *α*. The check indeed showed that the test has the expected false positive rate (Figure S4).

#### Comparison between asaMap and LAAA

To assess how asaMap performs compared to other similar methods, we performed a power comparison to the recently published LAAA test from Duan et al. (2017). The LAAA test is almost equivalent to comparing the M0 model to the M5 model in asaMap. Therefore, we performed the comparison by simply comparing the power of the M0 vs. M5 test and the M1 vs. M5 test in asaMap for all the 7 additive scenarios in Table 3. This showed that the M1 vs. M5 test always has slightly better power, because it has one less degree of freedom (Figure S4). It should be noted that the original LAAA test differs from the M0 vs M5 test in one way: it does not account for uncertainty in local ancestry inference and therefore only works when the ancestry information is known. Furthermore, the LAAA framework only has 1 proposed test. Hence besides being less powerful LAAA is also less flexible; asaMap also works with estimated admixture proportions and provides a range of different tests. For these reasons combined we did not perform more comparisons to LAAA.

### *TBC1D4* gene in a Greenlandic cohort

To further assess asaMap, we also applied it to real data from the Greenlandic population. This population is an Inuit population which is highly admixed: more than 80% of Greenlanders have some recent European ancestry and the Greenlanders have on average approximately 25% European ancestry (Moltke et al. 2015). A recent GWAS in the Greenlandic population (Moltke et al. 2014) led to the identification of a variant in the gene *TDB1D4* which confers high risk of type 2 diabetes and highly elevated 2-h plasma glucose levels. The lead SNP in the discovery part of this study was rs7330796 and to locate the causal variation four coding SNPs in high LD was identified using exome sequencing and subsequently genotyped. Among these SNPs rs61736969 located in *TBC1D4* was identified as the causative variant and shown to have a recessive effect.

Based on genotype and phenotype data from the Greenlandic IHIT cohort described in (Moltke et al. 2014), we tested the five above mentioned SNPs for ancestry-specific association with 2-h plasma glucose levels as a quantitative trait using a recessive model (for results see Figure 6). For the four non-causal SNPs; rs7330796 (original lead SNP), rs1062087, rs2297206, and rs77685055, we observed a more significant recessive effect of the variant when both alleles are inherited from the Inuit population (R4 vs. R7) compared to the standard GLM based test (R6 vs. R7), supporting our simulation based observation that asaMap can increase the power to detect associations when the causal SNPs remains untyped. Furthermore, for all four non-causal SNPs asaMap (R6 vs. R2) showed that the effect of carrying two effect alleles both inherited from the ancestral Inuit population is significantly different from the effect of carrying two effect alleles of which at least one is inherited from the ancestral European population. This suggests that these four SNPs are not causal. On the contrary, for rs61736969 the p-value for the population specific test for a recessive effect in the Greenlandic population is identical to the p-value of the standard test (R6 vs R7), which is consistent with the conclusion from (Moltke et al. 2014) that it is casual.

**Figure 6:**
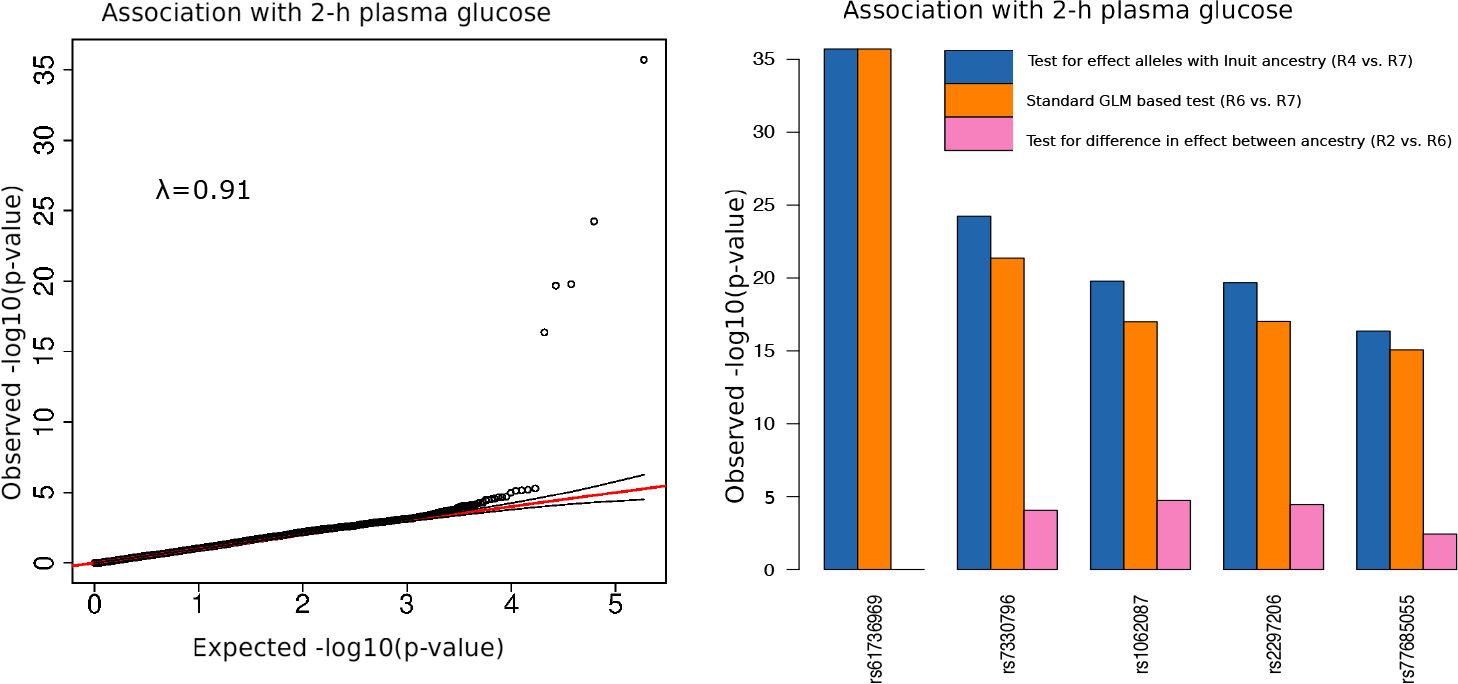
Association results for data from Greenland. Left: QQ-plot for 2-h plasma glucose in the Greenlandic IHIT cohort achieved using the asaMap recessive model for quantitative traits. Specifically, we tested for an effect in the ancestral Inuit population using the test R4 vs. R7. Right: Minus log10 of the p-values from three different statistical tests implemented in asaMap.

Finally, we note that the QQ-plot in Figure 6 shows that the ancestry-specific association test is not more inflated than regular association mapping tests, when using the first ten principal components to correct for population structure as described in Materials and Methods.

## Discussion

In this paper, we have presented asaMap, a flexible statistical testing framework for association mapping in admixed populations, which allows for the possibility that a tested variant can have different effects in the different ancestral populations. asaMap does this by modeling the local allelic ancestry as a latent variable.

Using simulated data we have demonstrated that asaMap provides ancestry-specific effect estimates that are unbiased and consistent. Furthermore, we have assessed how powerful asaMap’s association tests are compared to the standard GLM based tests, which are commonly used for performing association mapping in admixed populations. Unlike asaMap, these commonly used tests do not allow for the possibility that a tested variant can have different effects in the different ancestral populations. On the contrary, they are based on the assumption that the effect of the tested variant is the same regardless of its ancestry. This assumption is reasonable for a causal variant, but may not hold for the SNPs tested in a GWAS, which are usually not causal. Notably, we have here demonstrated that when the effect does depend on ancestry, the full test in asaMap, which tests if there is an effect in any population while allowing for ancestry-specific effects (M1 vs. M5) will outperform the commonly used GLM based tests. However, the gain in power depends strongly on the allele frequencies in the ancestral populations. If there is a lower frequency in the population in which the effect is highest than in the other population the gain in power can be very substantial. Conversely, if the frequency is higher, then gain in power can be negligible. Also our results suggest that the gain depends on the underlying ancestry distribution. Specifically, the power gain is largest when the ancestry specific effects are in the least contributing ancestral population, which makes sense because the alleles of the biggest ancestral population will reflect the alleles of the total population to a higher degree.

When the effect is not ancestry-specific, the full test in asaMap is less powerful than the commonly used test, which could be expected since the full test is based on a more complicated model. However, here the difference in power is small.

In addition to the full test, asaMap also allows for testing if a variant has an effect in a specific ancestral population - both with or without assuming that there is an effect in the other population. Testing if a variant has an effect in one of the ancestral populations, while allowing for an effect in the other population (M1 vs. M2 or M1 vs. M3) will be less powerful for identifying new associations than testing if a variant has an effect in one of the populations, while assuming no effect in the other population (M2 vs. M5 and M3 vs. M5). However, the former are useful tests for establishing if the causal variant is present and in LD with the tested variant in a specific population. Also, if one of the populations has already been extensively studied for a very large number of individuals, and no effect has been detected here, then it can be practical to assume that there is no effect in that population: our results show that the tests M2 vs. M5 and M3 vs. M5 are the most powerful if the effect is actually absent in one population. We therefore recommend using the tests M2 vs. M5 for association testing in datasets where very large-scale association tests have already been applied to one of the ancestral populations.

All the power results described above were based on simulations of variants with additive effects. For variants with recessive effects it is a bit more complicated since there are three possible effects for individuals carrying two effect alleles assuming there are two ancestral populations. However, we have here demonstrated that when an allele only has a recessive effect when both copies of it are inherited from one of the two populations there is potential to gain a great amount of power by allowing for ancestry-specific effects. More specifically, we observed a large gain in power when the tested variant was in high frequency in both populations, both for the full test (R1 vs. R7) and even more so for the test for an effect in a specific ancestral population (R4 vs R7). We expect the same to be true in all cases where the frequency of the effect allele is high in the population where it does not have an effect, because in this case a lot of the individuals carrying two copies of the effect allele will not be affected, causing the standard GLM based recessive association test to have low statistical power.

Another useful test is M1 vs. M4, which tests whether the effect sizes are different in the two ancestral populations. Since we expect the effect of a causal allele to be similar in the two ancestral populations a significant test is an indication that a variant is not causal. Two fairly different populations can have different amounts of LD between the causal site and the tested variant, but this may not always be true, thus a non-significant test for different effect sizes can clearly not be taken as evidence that the variant is causal. We realize that expecting the causal allele to have similar effects in the ancestral population suggests that the gain in power from using asaMap may be limited in an ideal GWAS study, where all potentially causal variants have been well imputed from large, deeply sequenced reference panels. However, currently most GWAS in admixed populations deviate from this ideal in that imputation may be challenging or that reference panels may be limited. Specifically, large, deeply sequenced reference panels that adequately represent potential founder events in one of the ancestral populations may not be available.

Finally, asaMap offers a test for whether the assumed non-effect allele, i.e. the allele that is not assumed to be the effect allele in the default asaMap model, has an ancestry-specific effect. The motivation for this is that asaMap, just like other similar methods (Pasaniuc et al. 2011, Yorgov et al. 2014), focuses its tests on the potentially different effects of a specific assumed effect allele. However, it could in principle be the other allele that has an ancestry specific effect. If this is the case it will not lead to false positives, but it may result in suboptimal power to detect an effect and particularly to detect difference in effects between populations. We have not experienced this as a problem in practice, but since it may occur, offering this additional test seems potentially useful to the users.

asaMap allows local ancestry probabilities from one of the following two sources. By default, asaMap calculates local ancestry probabilities based on individual genome-wide admixture proportions and ancestral allele frequencies. Alternatively, local ancestry probabilities can be provided by the user. In addition to the power gained by asaMap using the default calculation of local ancestry probabilities from admixture proportions, we demonstrate a hypothetical gain in power from working with true (but unobservable) local ancestry. Thus, if accurate local ancestry inference is possible we encourage the user to provide local ancestry probabilities to asaMap. However, if the users observe high uncertainty in the local ancestry inference i.e. low probabilities (<95%), we recommend providing individual genome-wide admixture proportions and ancestral allele frequencies to asaMap and running it using the default setting.

To test asaMap on real data, we applied it to genotype data from admixed individuals from Greenland for five SNPs and corresponding levels of 2-h plasma glucose. We demonstrated that the ancestry-specific tests in asaMap can increase the statistical power of the GWAS when the causal variant remains untyped. Also, asaMap correctly provided results that were consistent with the identified causal loss of function variant being causal. Furthermore, asaMap correctly provided results, which support that the four remaining SNPs have ancestry-specific effects and thus are unlikely to be causal.

In summary, we have shown, using both simulated and real data, that asaMap by allowing for ancestry-specific effects provides tests that in some cases are much more powerful than the standard GLM based tests that are commonly used in GWAS. We have also shown the asaMap tests, at least the full test, are almost as powerful as the standard GLM based test in all other investigated cases. Finally, we have shown that asaMap can be used to test if a variant has an ancestry dependent effect, which can be helpful for assessing if a tested SNP is causal. This suggests that asaMap is a powerful and flexible complement to the standard tests commonly used when carrying out a GWAS in admixed populations.

## Supporting Information

### Supplementary figures

**Figure S1:**
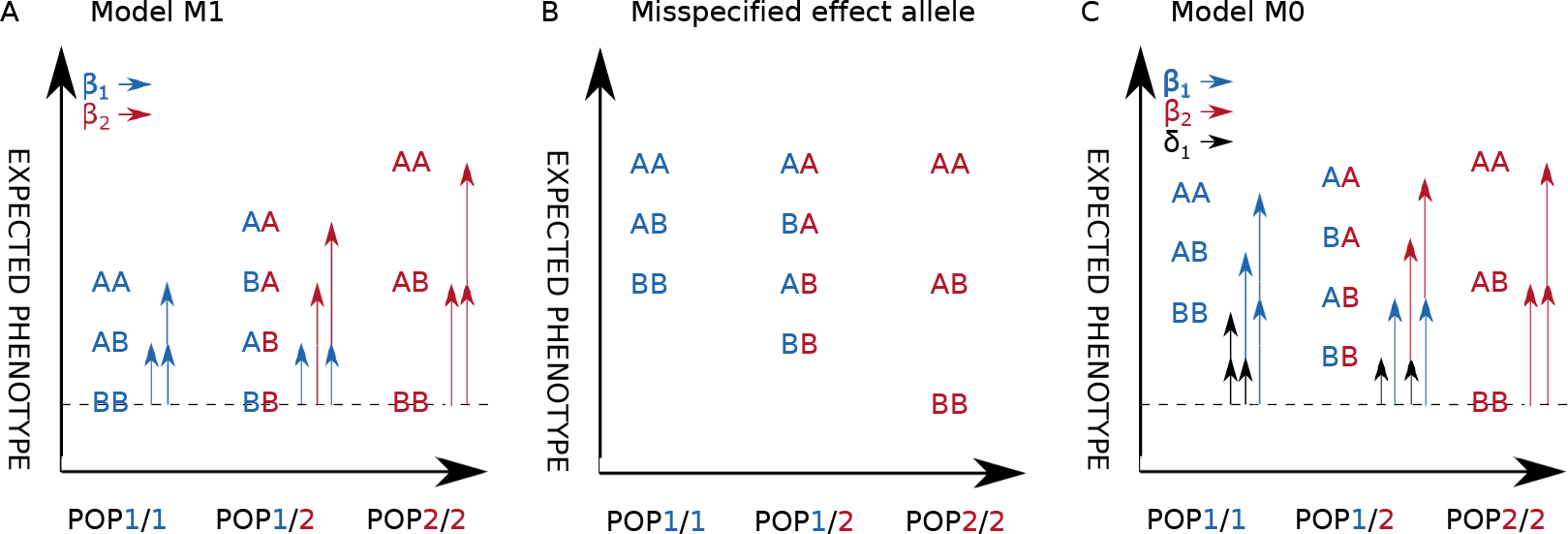
Population-specific effect of the assumed non-effect allele. A: Model M1 allows ancestry-specific effect of allele A and thus implicitly assumes that the expected phenotype for BB individuals does not depend on the local ancestry. B: If allele B mediates the population-specific effect, the expected phenotype for BB individuals depend on the local ancestry and allele A is misspecified as the assumed effect allele. C: Model M0 allows an effect of the assumed non-effect allele B, when inherited from ancestral population 1.

**Figure S2:**
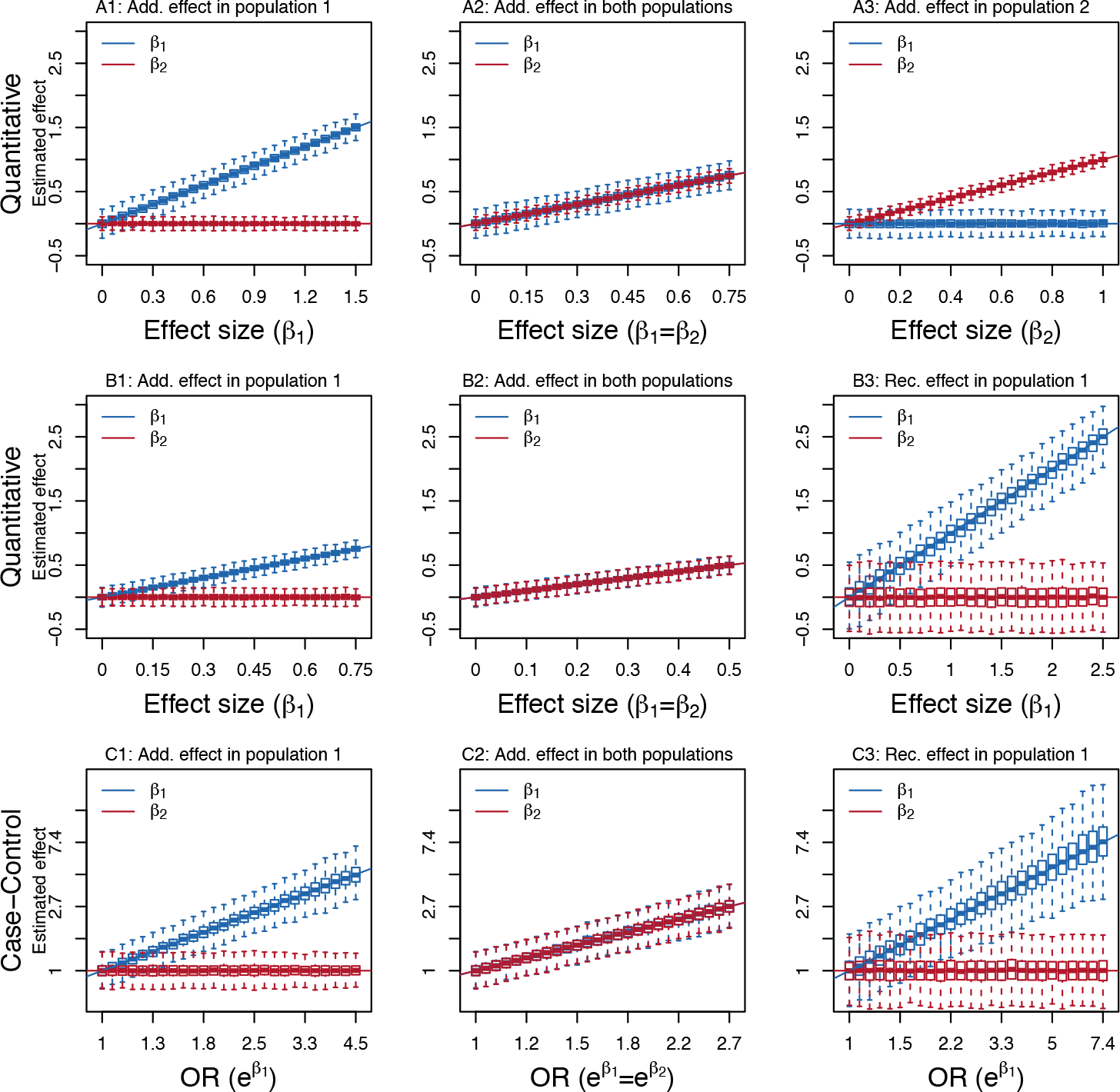
Results of bias simulations for all nine scenarios (A1-C3). Each box show the results of 1000 simulations. The lines show the true effect sizes. *β*_1_ and *β*_2_ denote the effect sizes in population 1 and 2, respectively. The simulated scenarios are described in Table 3 and the tests are described in Table 2 and Table S2.

**Figure S3:**
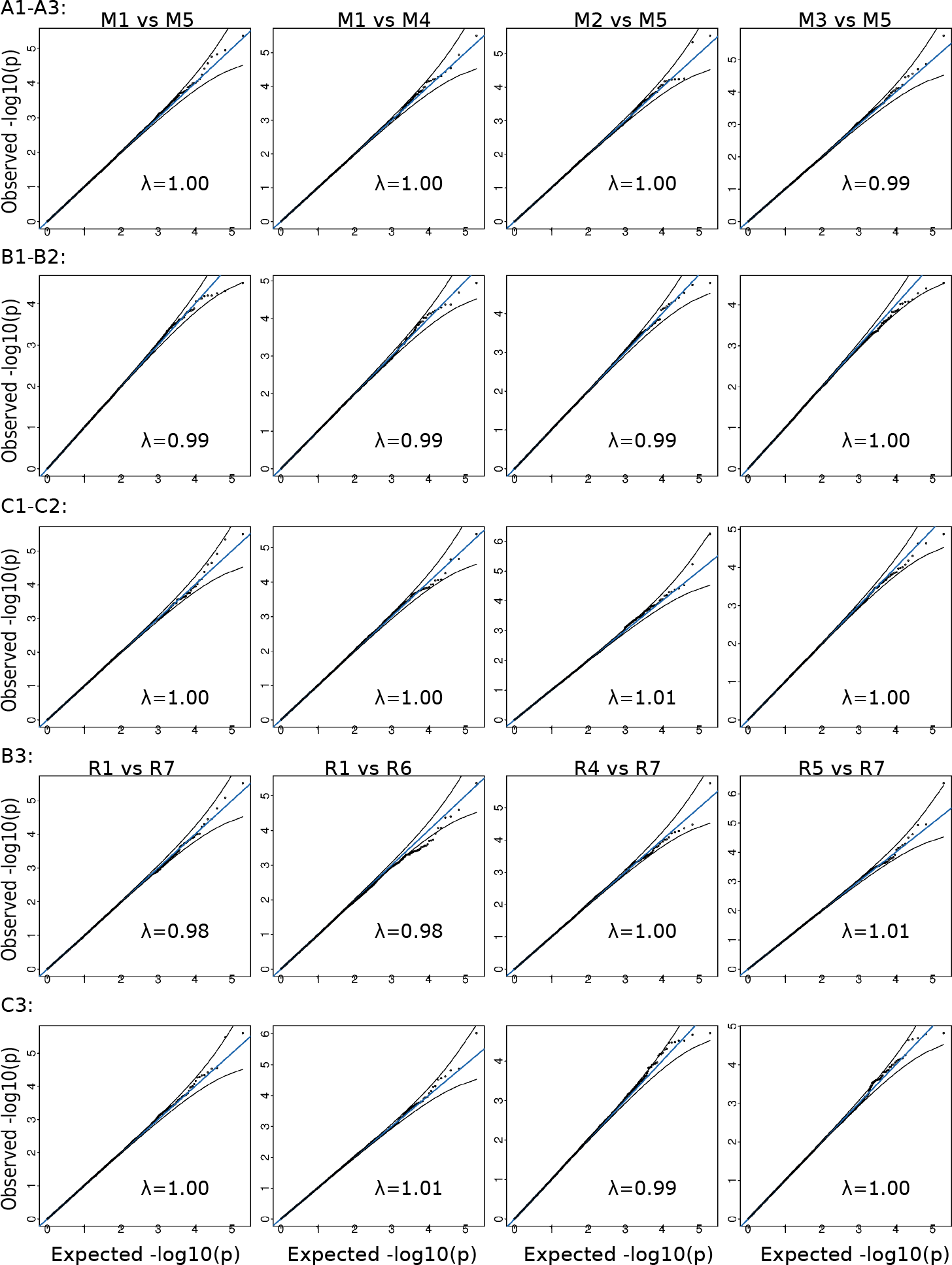
QQ-plots of the p-values obtained by applying asaMap to simulations from the null model of all simulation scenarios (A1-C3). Each QQ-plot is based on 100000 simulations. The simulation scenarios are described in Table 3 and the tests are described in Table 2 and Table S2. The three top rows show results for the asaMap tests that assume an additive genetic effect, while the two bottom rows show results for the asaMap tests that assume a recessive genetic effect. The genomic inflation factor λ is stated for each scenario.

**Figure S4:**
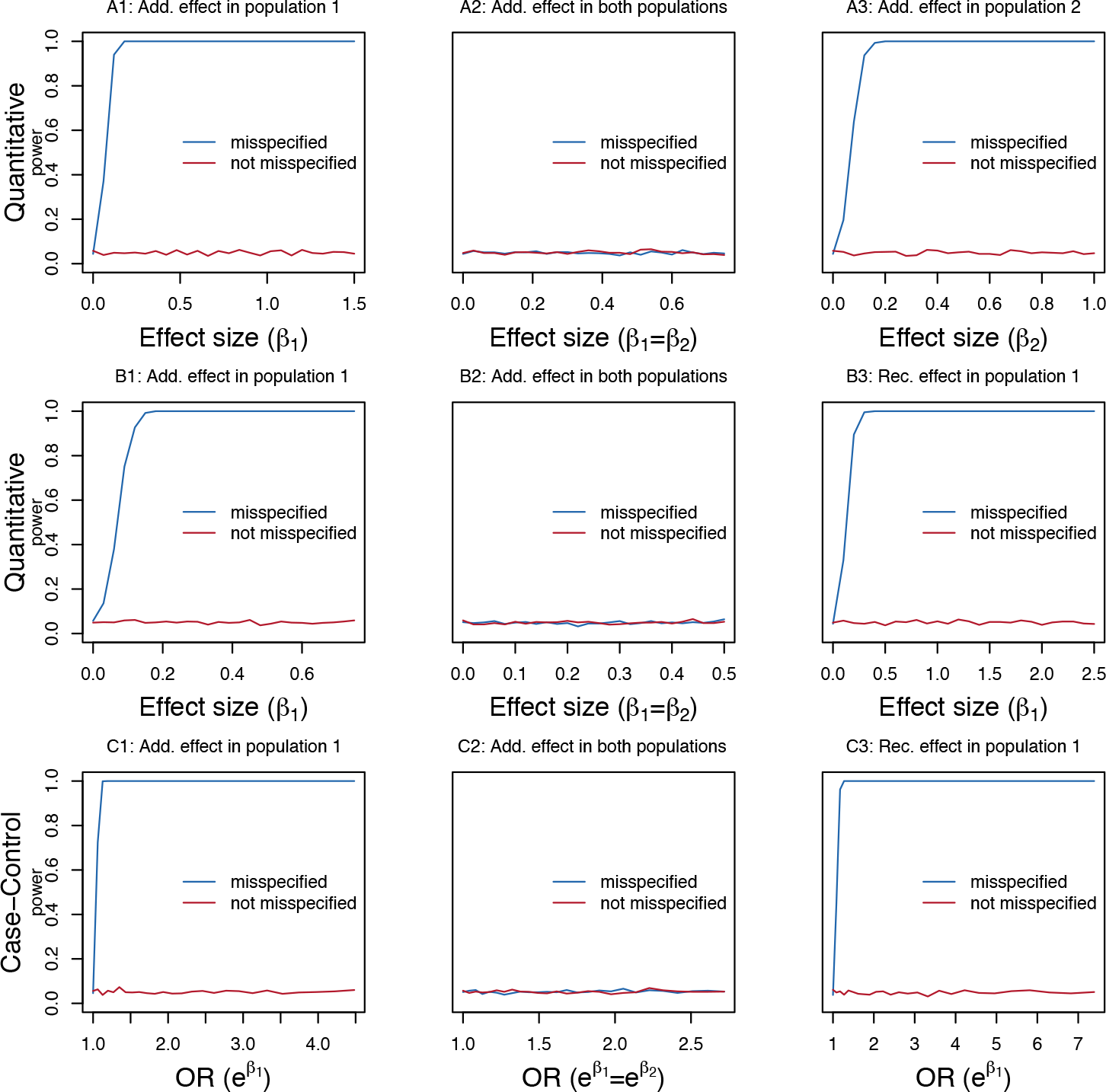
Results for the tests of whether the assumed non-effect allele has an effect applied to simulated data from nine scenarios (A1-C3). The simulation scenarios are described in Table 3. The tests are based on comparing M0 vs. M1 and R0 vs. R1. M0 and R0 are described in equations 4 and 7. All the curves are based on 1000 simulations for each effect size for each scenario and show the fraction of the tests that led to p-values that are smaller than 0.05. The curves labeled “misspecified” show the results of applying the tests with the assumed effect allele misspecified. The curves labeled “not misspecified” show the results of applying the tests with the assumed effect allele correctly specified.

**Figure S5:**
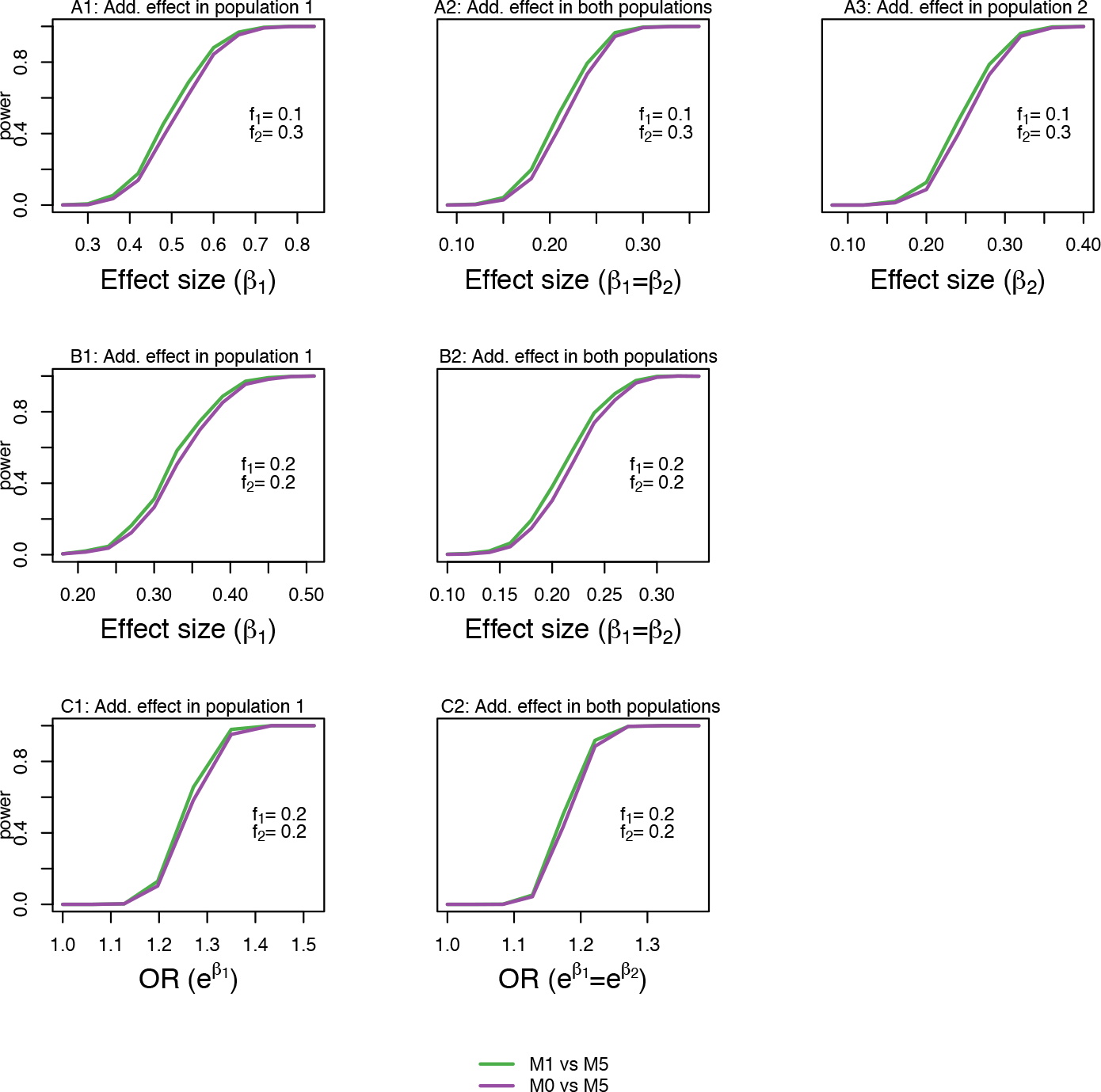
Results for comparing statistical power for tests M1 vs. M5 and M0 vs. M5. M0 vs. M5 is the same test as proposed in Duan et al. (2017) where it is named “LAAA”. Both tests have been applied to simulated data from the 7 additive scenarios from Table 3. The tests are based on comparing M1 vs. M5 and M0 vs. M5. M0 is described in equations 4. All the curves are based on 1000 simulations for each effect size for each scenario and show the fraction of the tests that led to p-values that are smaller than 10^−8^. The curves labeled M0 vs. M5 show the results of applying the proposed LAAA test (Duan et al. 2017).

**Figure S6:**
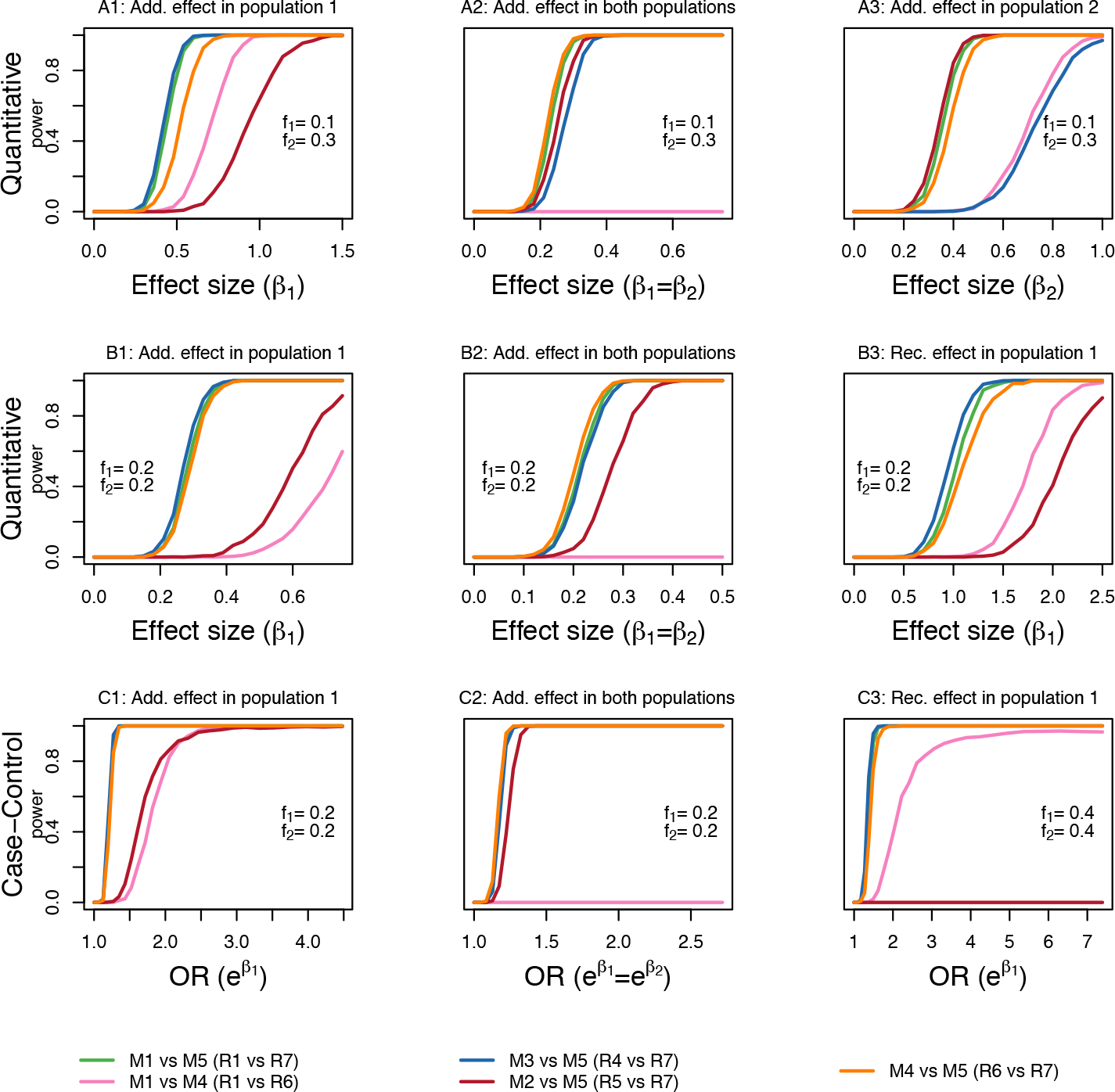
Same simulations as in Figure 3. The admixture proportions are not as stated in Table 3, rather they are sampled from admixture proportions in a cohort of 4629 Greenlanders, estimated using ADMIXTURE. This means that population 1 is now the bigger population, in terms of the amount of DNA.

**Figure S7:**
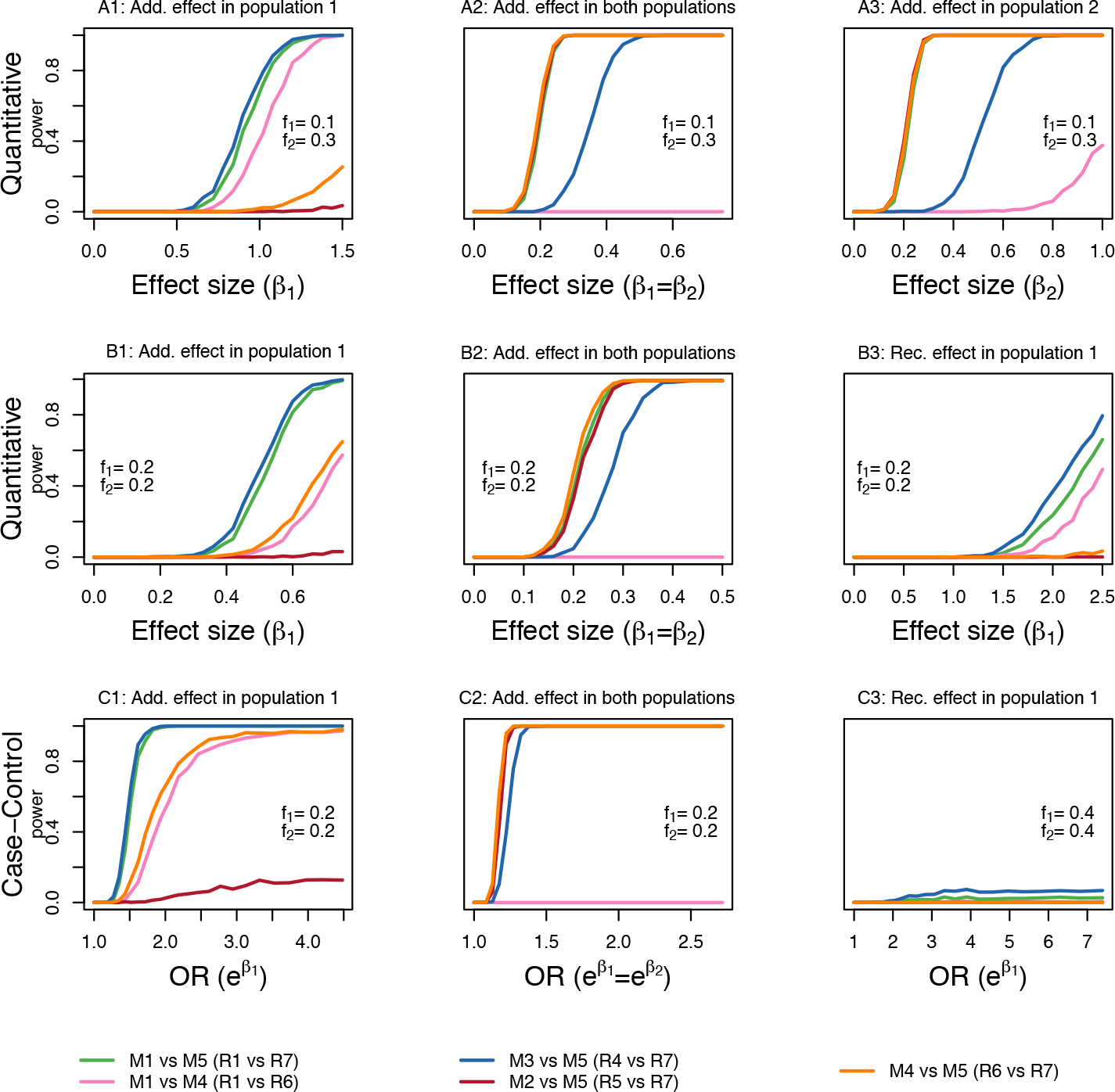
Same simulations as in Figure S6. Here the admixture proportions for population 1 and 2 have been swapped around. This means that population 1 is now the smaller population, in terms of the amount of DNA.

### Supplementary tables

**Table S1:** Recessive ancestry-specific genetic effects models. *β*_1_ is the effect of carrying two copies of the assumed effect allele when both are inherited from population 1, *β*_2_ is the effect carrying two copies of the effect allele when both are inherited from population 2 and *β*_*m*_ is the effect of carrying two copies of the effect allele when one of the copies is inherited from population 1 and the other from population 2.

| Model | Hypothesis | The model assumes |
| --- | --- | --- |
| R1 | $(\beta_1, \beta_m, \beta_2) \in R^3$ | Ancestry-specific effects |
| R2 | $\beta_1 \in R, \beta_m = \beta_2 \in R$ | Same effect when 1 or both effect alleles are from population 2 |
| R3 | $\beta_1 = \beta_m \in R, \beta_2 \in R$ | Same effect when 1 or both effect alleles are from population 1 |
| R4 | $\beta_1 \in R, \beta_m = \beta_2 = 0$ | Only an effect when both effect alleles are from population 1 |
| R5 | $\beta_1 = \beta_m = 0, \beta_2 \in R$ | Only an effect when both effect alleles are from population 2 |
| R6 | $\beta_1 = \beta_m = \beta_2 \in R$ | Same effect regardless of ancestry |
| R7 | $\beta_1 = \beta_m = \beta_2 = 0$ | No effect |

**Table S2:** Relevant tests comparing the different nested recessive ancestry-specific genetic effects models described in Table S1.

| Models | Tests if there is |
| --- | --- |
| R1 vs. R7 | A recessive effect for some combination of ancestry |
| R1 vs. R4 | Only an effect when both effect alleles are from population 1 |
| R1 vs. R5 | Only an effect when both effect alleles are from population 2 |
| R1 vs. R6 | Any ancestry dependence of the effect |
| R2 vs. R6 | A different effect when both effect alleles are from population 1 |
| R3 vs. R6 | A different effect when both effect alleles are from population 2 |
| R4 vs. R7 | A recessive effect in population 1 |
| R5 vs. R7 | A recessive effect in population 2 |
| R6 vs. R7 | A recessive effect assuming it is independent of ancestry |

### Supplementary text

Below we describe the models and methods that were not described in detail in the main text.

#### Case-control study modification

For case-control studies or other dichotomous traits studies asaMap is based on a logistic regression analysis, since this allows the inclusion of additional covariates (such as principal components) in the model. Specifically, given the locus state *s*, the probability *π*_*i*_ that individual *i* is affected is modeled by

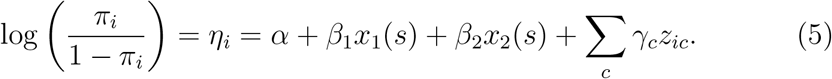

where *x*_1_(*s*) and *x*_2_(*s*) are the design matrix entries for the two ancestry-specific effects and *z*_*ic*_ is the value of covariate *c* for individual *i*.

#### Overview of recessive models and tests

In addition to the models that assume an additive genetic effect, models that assume a recessive effect have also been implemented in asaMap. Their design is slightly more complicated than the additive models. This is because we wish to have the best possible power to detect the genetic effect of a recessive disease causing variant that may remain untyped and only be present in one of the ancestral populations. Notably, due to this more complex modelling, the recessive genetic model is complicated for more than two ancestral populations.

A recessive model in general assumes that there only is an effect, when an individual carries two copies of the effect allele. The full ancestry-specific recessive model, R1, includes three different effects: *β*_1_ is the effect of carrying two copies of the effect allele when both are inherited from population 1, *β*_2_ is the effect of carrying two copies of the effect allele when both are inherited from population 2 and *β*_*m*_ is the effect of carrying two copies of the effect allele when one of the copies is inherited from population 1 and the other from population 2. The linear predictor for this model is given by

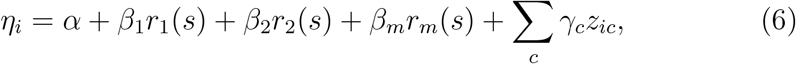

where *r*_1_ = 1 if the individual carries two copies of the effect allele that are both inherited from population 1 and *r*_1_ = 0 otherwise, *r*_2_ = 1 if the individual carries two copies of the effect allele that are both inherited from population 2 and *r*_2_ = 0 otherwise and where *r*_*m*_ = 1 if the individual carries two copies of the assumed effect-allele and has inherited one copy from each of population 1 and 2.

This allows us to fit both R1 and a range of sub-models (see Table S1 for an overview and later text for a more detailed description). The models R4 and R5 are of most interest. In R4 it is assumed that there is no effect of carrying two copies of the effect allele - unless both alleles are inherited from population 1. This model can then be compared against R7 - where it is assumed that there is no effect - to test if there is a significant recessive effect of the effect allele when inherited from population 1. To test if the model assumption for model R4 is appropriate we can compare against R1 and to test if there is a different recessive effect when both alleles are inherited from population 1 than otherwise, we can compare R2 to R6, where the effect is assumed to be independent of ancestry. In the exact same way R5 vs. R7 can be used to test for a recessive effect when both variants are inherited from population 2 and R1 vs. R5 can be used to check the assumptions of this test.

We have also implemented an expanded model, R0, which compared to model R1 includes two additional effects: the effect of carrying two copies of the assumed non-effect allele that are both inherited from population 1 and the effect of carrying two copies of the assumed non-effect allele that are both inherited from population 2. The model is defined as follows:

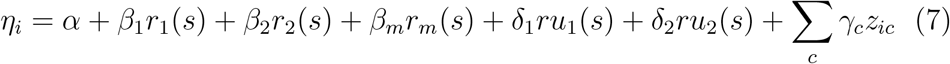

where, *r*_1_, *r*_2_, *r*_*m*_, *β*_1_, *β*_2_ and *β*_*m*_ are defined as above, *ru*_1_ is an indicator of whether an individual carries two copies of the assumed non-effect allele that are both inherited from population 1, *ru*_2_ is an indicator of whether an individual carries two copies of the assumed non-effect allele that are both inherited from population 2 and *δ*_1_, *δ*_2_ are the effect sizes corresponding to *ru*_1_ and *ru*_2_. By including these additional effects the model allows the user to check if the assumed effect allele is misspecified. This check is performed by testing model R0 against model R1, to see if we reject the null hypothesis of *δ*_1_ = *δ*_2_=0. Note that, for this we assume that carrying two copies of the assumed non-effect allele with two different ancestries will not have an effect that *δ*_1_ or *δ*_2_ does not detect.

#### Detailed equations for the additive models

Below we provide a more detailed description of the different nested models used under the assumption of an additive genetic effect outlined in Equation 2, Equation 4 and Table 1. More specifically, we list the linear predictors for all the models. In this list, the covariates *x*_1_(*s*) and *x*_2_(*s*) denote the counts of the assumed effect allele from population 1 and 2, respectively for a locus in locus state *s*. In addition, *α* parameterizes the intercept (baseline) and additional covariates enters the model in the *z*_*ic*_ with effects γ_*c*_. Finally, we use *δ*_1_ to parameterize the effect of the assumed non-effect allele and *u*_1_ is the observed number of assumed non-effect alleles from population 1. With this notations the linear predictors are:

**M0:**

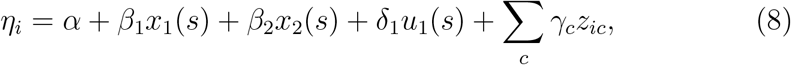
**M1:**

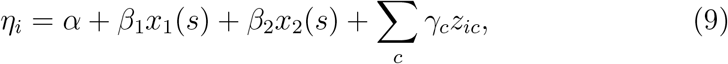
**M2:**

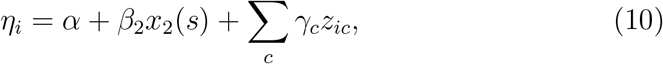
**M3:**

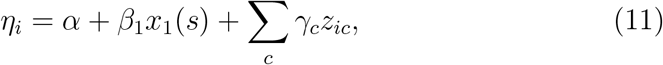
**M4:**

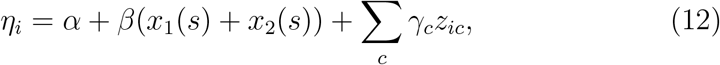
**M5:**

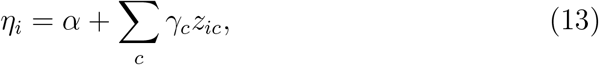

#### Detailed equations for the recessive models

Below we provide a more detailed description of the different nested models used under the assumption of a recessive genetic effect outlined in Equation 6, Equation 7 and Table S1. More specifically, we list the linear predictors for all the models.

In this list, *r*_1_ and *r*_2_ indicates if the individual carries two copies of the assumed effect allele and both are inherited from population 1 or 2, respectively. Similarly, *r*_*m*_ indicates if the individual carries two copies of the assumed effect allele and they are inherited from both populations. The corresponding effects are denoted by *β*_1_, *β*_2_ and *β*_*m*_. In addition, *α* parameterizes the intercept (baseline) and additional covariates enters the model in the *z*_*ic*_ with effects γ_*c*_. Finally, *ru*_1_, *ru*_2_ indicates if the individual carries two copies of the assumed non-effect allele and both alleles are inherited from population 1 or 2, respectively and *δ*_1_, *δ*_2_ are the corresponding effects. With this notation the predictors are:

**R0:**

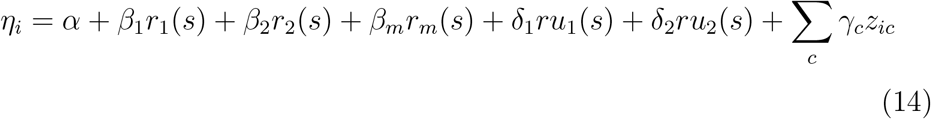
**R1:**

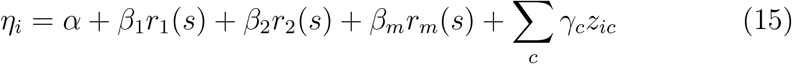
**R2:**

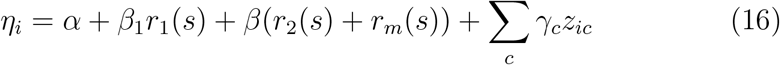
**R3:**

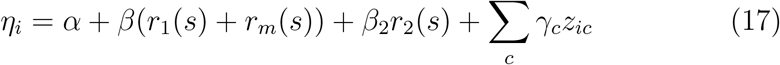
**R4:**

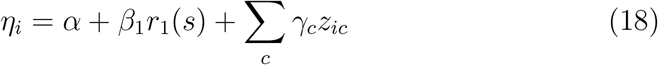
**R5:**

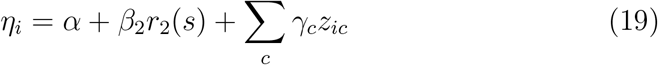
**R6:**

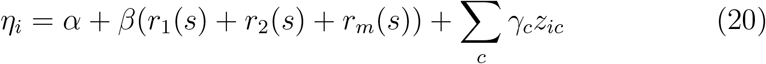
**R7:**

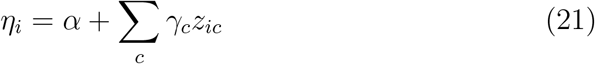

#### EM algorithm

Below we will go through the details of the EM algorithm used in asaMap.

##### Notation

We will use the following notation:

*n* individuals
*y*_*i*_ phenotype of ind *i* ∈ {1, …, *N*}
*g*_*i*_ genotype of ind *i*, *g* ∈ {0, 1, 2}
*Q*_*i*_ admixture proportions, for individual *i*. For convenience instead of *Q*_*i*_ we also use the notation *q* = {*q*_1_, *q*_2_} for the two admixture proportions.
*f* = {*f*_1_, *f*_2_} population specific allele frequencies
*ϕ* vector of effect sizes e.g. (*β*_1_, *β*_2_) and other regression parameters e.g. (*α*, γ, *σ*)
*s* locus state, *s* = {*a*, *t*} where *a* = (*a*_1_, *a*_2_) is the ordered information on ancestry and *t* = (*t*_1_, *t*_2_) is the ordered allelic genotype.

##### Likelihood functions

The likelihood for the observed data, assuming that individuals are independent given genotypes, admixture proportions and population specific allele frequencies, is given by:

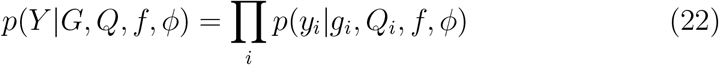

and splitting the probabilities according to locus state (see Figure 2) gives

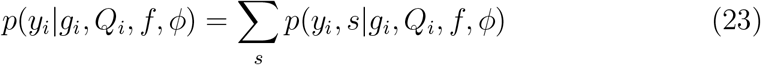

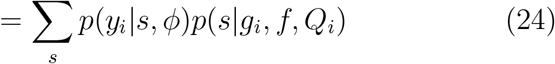

Conditional on locus state, the phenotype follows a normal distribution.

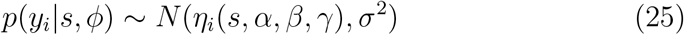

with

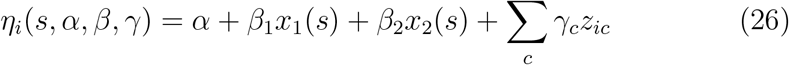

where we use that for population *k* we have 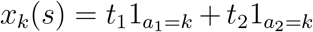 for the additive genetic model. The normal distribution is part of the exponential family. The density of a normal with mean *η* and standard deviation *σ* is

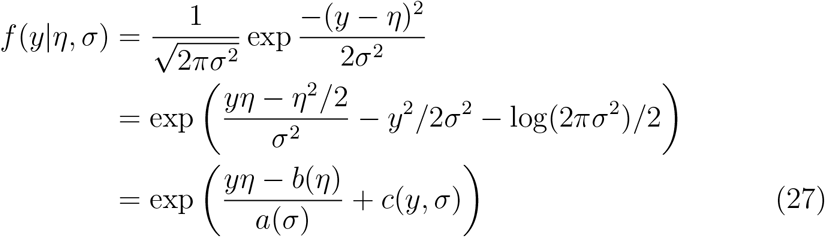

with *b*(*η*) = *η*^2^/2, *a*(*σ*) = *σ*^2^ and *c*(*y*, *σ*) = −*y*^2^/2*σ*^2^ − log(2*πσ*^2^)/2.

##### Derivation of EM algorithm

The expression that must be maximized in a single EM algorithm step is:

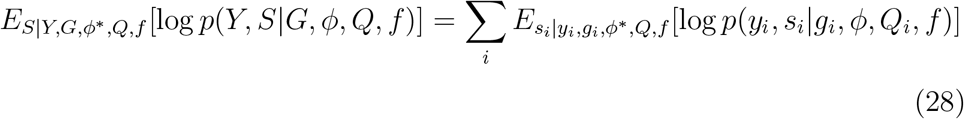

as a function of all regression parameters *ϕ*, where *ϕ*\* is fixed to the parameter values from previous iteration. Using *p*(*y*_*i*_, *s*_*i*_|*G*, *ϕ*, *Q*_*i*_, *f*) = *p*(*y*_*i*_|*s*_*i*_, *ϕ*)*p*(*s*_*i*_|*Q*_*i*_, *f*) where the second term does not depend on the parameters to be optimized, this is equivalent to maximizing

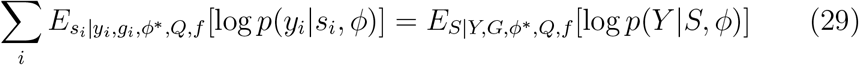

Following (Lake et al. 2003) we take the derivative with respect to the vector of ancestry-specific effect sizes and get:

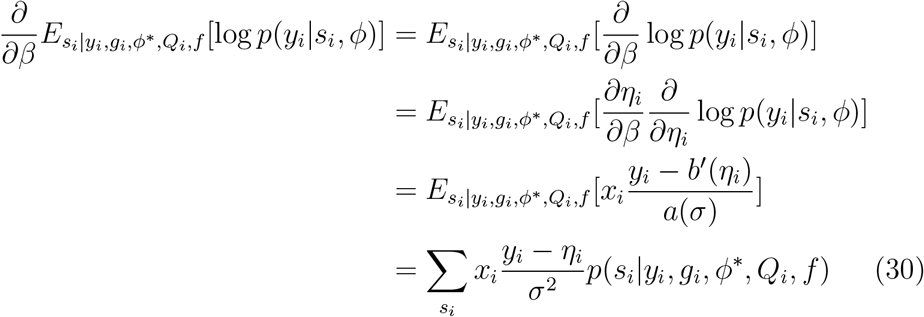

where *x*_*i*_ = (*x*_1_(*s*_*i*_), *x*_2_(*s*_*i*_)) and *η*_*i*_ = *η*(*x*_*i*_, *z*_*i*_, *α*, *β*, γ). The equivalent formula holds for the derivative with respect to *α* and γ. We therefore get

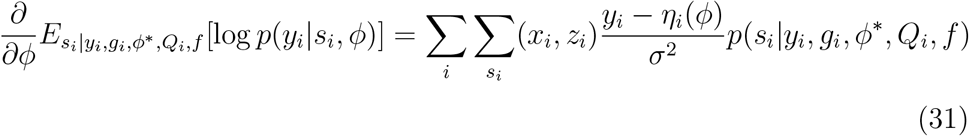

which is recognized as the score function for a weighted regression where each individual, *i*, contributes one observation per possible state *s*_*i*_ and where the weights, *p*(*s*_*i*_|*y*_*i*_, *g*_*i*_, *ϕ*\*, *Q*_*i*_, *f*), are the posterior distribution of states given all the observed data and based on the previously fitted parameters (see below for details). The same formula holds for the logistic regression. The updated regression parameters *ϕ* can therefore be estimated by fitting a weighted linear regression in case of a quantitative trait and a weighted logistic regression for case/control data.

##### Posterior distribution of locus state

The conditional distribution of locus state given previous parameters, observed data and genotype is found using

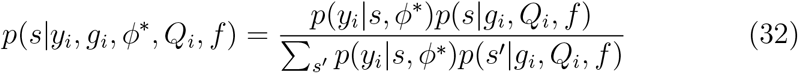

where *p*(*s*|*g*_*i*_, *Q*_*i*_, *f*) is given in equation 3 and *p*(*y*_*i*_|*s*, *ϕ*\*) is the phenotype distribution given locus state and previous parameters.

##### Optimization strategy for normal distributed trait

First a rough initial guess of the standard deviation is calculated by

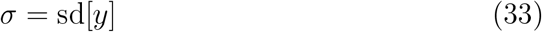

and randomly chosen start values for the regression parameters are sampled from a uniform distribution e.g.

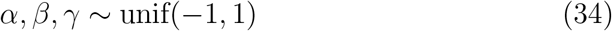

Then regression weights are calculated according to (32) and a weighted regression according to the score function in (31) is carried out to update *β, γ*. This is followed by an update of *σ* using the weighted sum of squared residuals from the weighted regression and *n*−*p* df, where *n* is the number of individuals and *p* is the number of effect parameters in the linear predictor e.g. (*p* = 3 + *C*) for M1 model with *C* being the number of γ parameters.

